# *Rice Yellow Mottle Virus* resistance by genome editing of the *Oryza sativa* L. ssp. japonica nucleoporin gene *OsCPR5.1* but not *OsCPR5.2*

**DOI:** 10.1101/2023.01.13.523077

**Authors:** Yugander Arra, Florence Auguy, Melissa Stiebner, Sophie Chéron, Michael M. Wudick, Manuel Miras, Van Schepler-Luu, Sébastien Cunnac, Wolf B. Frommer, Laurence Albar

**Author notes:** Equal contribution (Yugander Arra, Florence Auguy).

## Abstract

Rice yellow mottle virus (RYMV) causes one of the most devastating rice diseases in Africa. Management of RYMV is challenging. Genetic resistance provides the most effective and environment-friendly control. The recessive resistance locus *rymv2* (*OsCPR5*.*1*) had been identified in African rice (*O. glaberrima*), however, introgression into *O. sativa ssp. japonica* and *indica* remains challenging due to crossing barriers. Here, we evaluated whether CRISPR/Cas9 genome editing of the two rice nucleoporin paralogs *OsCPR5*.*1* (*RYMV2*) and *OsCPR5*.*2* can be used to introduce RYMV resistance into the *japonica* variety Kitaake. Both paralogs had been shown to complement the defects of the Arabidopsis *atcpr5* mutant, indicating partial redundancy. Despite striking sequence and structural similarities between the two paralogs, only o*scpr5*.*1* loss-of-function mutants were fully resistant, while loss-of-function *oscpr5*.*2* mutants remained susceptible, intimating that *OsCPR5*.*1* plays a specific role in RYMV susceptibility. Notably, edited lines with short in-frame deletions or replacements in the N-terminal domain (predicted to be unstructured) of *OsCPR5*.*1* were hypersusceptible to RYMV. In contrast to mutations in the single Arabidopsis *AtCPR5* gene, which caused severely dwarfed plants, *oscpr5*.*1* and *oscpr5*.*2* single *knockout* mutants show neither substantial growth defects nor symptoms indicative of programmed cell death, possibly reflecting functional redundancy of the isoforms regarding other important functions. The specific editing of *OsCPR5*.*1*, while maintaining *OsCPR5*.*2* activity, provides a promising strategy for generating RYMV-resistance in elite *Oryza sativa* lines as well as for effective stacking with other RYMV resistance genes or other traits.

## Introduction

Rice is one of the main staple food crops in developing countries, especially in sub-Saharan Africa (sSA). About 60% of the rice consumed is produced by sSA. Biotic and abiotic constraints impact rice yield in both irrigated and rainfed rice cultivation environments (Diagne *et al*., 2013). In sSA, rice yellow mottle virus (RYMV, genus *Sobemovirus*, family *Solemoviridae*) causes substantial losses and is often considered as the dominant rice disease in irrigated and lowland ecologies (Ochola *et al*., 2015; Suvi *et al*., 2020; Traoré *et al*., 2015). RYMV disease was first identified in East Africa in 1966 (Bakker, 1970) and has since been observed in almost all rice producing African countries. Typical symptoms are severe leaf mottling, yellow-green streaking, stunting at the vegetative stage, and reduced panicles emergence and sterility at the reproductive stage (Bakker, 1970). Depending on the environmental conditions and the susceptibility of the particular rice cultivars, in severe cases yield loss can exceed 50%. RYMV is transmitted *via* animal vectors - mammals and insects, as well as by inadequate agricultural practices, such as transplanting (Traoré *et al*., 2010). Prophylactic practices can be implemented to eliminate reservoirs, reduce infection at seedling stages and improve plant health, but their effect is limited and the most effective way to sustainably reduce the impact of the disease is the use of resistant varieties.

Varietal resistance sources were found in cultivated species. High resistance has been observed in about 7% of accessions of *O. glaberrima*, a domesticated African rice species, but is very rare in *O. sativa*, which is by far the most cultivated rice species around the world and in Africa (Pidon *et al*., 2020; Thiémélé *et al*., 2010). Three major resistance genes have been identified to date: *RYMV1 (eIF(iso)4G1), RYMV2* (*CPR5*.*1*), and *RYMV3* (NB-LRR) (Albar *et al*., 2006; Bonnamy *et al*., 2023; Odongo *et al*., 2021; Orjuela *et al*., 2013; Pidon *et al*., 2017). Interspecific sterility barriers hamper the introgression of interesting traits from *O. glaberrima* to *O. sativa* (Garavito *et al*., 2010). Thus breeding for RYMV resistance had focused primarily on *rymv1-2*, the only resistance allele originating from *O. sativa* (Bouet *et al*., 2013; Jaw *et al*., 2012). *RYMV2*-mediated resistance is inherited recessively; six alleles have been described in *O. glaberrima* (Pidon *et al*., 2020). Notably, the resistance in all these alleles is caused either by frameshift mutations or premature stop codons, leading to truncated proteins (7 to 83%), and thus loss of function.

*RYMV2* (gene name *OsCPR5*.*1* in *O. sativa*) is a close homolog of Arabidopsis *AtCPR5* (constitutive expresser of Pathogenesis Related genes 5) (Orjuela *et al*., 2013; Pidon *et al*., 2020). AtCPR5 is a membrane protein that functions as an integral membrane nucleoporin in transport of cargo across the nuclear pore (Gu *et al*., 2016; Xu *et al*., 2021). *atcpr5* mutants show broad spectrum resistance against bacterial, fungal and oomycete pathogens, however, they show severe growth defects which are consistent with a central role as part of the nucleopore complex. The exportin *XPO4* was identified as a genetic interactor of *CPR5* (Xu *et al*., 2021). Moreover, ROS levels and/or signaling (Jing and Dijkwel, 2008), unfolded protein response (UPR) (Meng *et al*., 2017), endoreduplication, cell division, cell expansion and spontaneous cell death were affected in *atcpr5* mutants (Kirik *et al*., 2001; Perazza *et al*., 2011). However, due to the severe defects such as dwarfing, the gene is not considered suitable for biotechnological applications (Bowling *et al*., 1997). The mechanism by which loss-of-function in *oscpr5*.*1* (*rymv2*) causes RYMV resistance is unknown. While the *Arabidopsis* genome contains only a single *AtCPR5* copy, two paralogs were found in other species, including rice (*OsCPR5*.*1; OsCPR5*.*2*).

Marker assisted backcross breeding (MABB) is widely used to introgress important major resistance genes into susceptible varieties (Yugander *et al*., 2018). However, in both conventional or MABB, introgression of single or multiple resistance genes into an elite variety is time-consuming and may not be able to remove unwanted linked loci and traits, even after multiple backcrosses, due to linkage drag. Introgression of resistance genes from *O. glaberrima* into *O. sativa* varieties is hampered by the crossing barrier (Heuer and Miézan, 2003). To overcome these constraints, innovative plant breeding technologies, particularly the clustered regularly interspaced short palindromic repeats (CRISPR) associated (Cas9) system have been used successfully in field crops including rice (Endo *et al*., 2016; Jiang *et al*., 2013; Oliva *et al*., 2019; Zhang *et al*., 2014). To evaluate whether editing can be used to effectively generate new *RYMV2* resistance alleles in *O. sativa*, and to analyze the role of the two *OsCPR5* paralogs in resistance, CRISPR/Cas9 was used to edit the two genes in *Oryza sativa ssp. japonica* cv. Kitaake. Loss-of-function mutants in both genes and mutants with short in frame mutations in *OsCPR5*.*1* were evaluated. We observed effective resistance only in *oscpr5*.*1* loss-of-function mutants, but not in *oscpr5*.*2* loss-of-function mutants. By contrast to the *Arabidopsis atcpr5* mutants, *oscpr5*.*1* and *oscpr5*.*2* knockout mutants did not show detectable defects under greenhouse conditions. We conclude that *OsCPR5*.*1* and *OsCPR5*.*2* have distinct roles in virus resistance and may have overlapping functions regarding other physiological functions. Our data indicate that editing provides an option to stack resistance genes against RYMV to better protect against virus strains that overcome resistance, and to stack with other disease resistance genes (Bonnamy *et al*., 2023; Hébrard *et al*., 2018; Pinel-Galzi *et al*., 2016).

## Results

### Bioinformatic analysis of *OsCPR5*.*1* and *OsCPR5*.*2*

To evaluate whether *OsCPR5*.*1* and *OsCPR5*.*2* might be redundant, or have distinct functions, we analyzed transcript levels in public RNAseq databases (Fig. S1). Both genes showed broad and overlapping expression across different tissues and developmental stages, and no major effects of various stresses on mRNA levels were found (Rice RNA-seq database, Zhailab@SUSTech, not shown). Structure predictions indicated that OsCPR5.1 and OsCPR5.2 had similar structures with a helix bundle from which a long helix extended, as well as possibly unstructured domains at the N-terminus (Fig. 1a). Prediction of homo- and heterodimer structures is shown in (Fig. S2); however, the confidence level for the dimer interface is low. For all three CPR proteins, similar dimeric structures were found among the highest ranked models, only the heterodimer of OsCPR5.1 and OsCPR5.2 appeared different; however, this does not provide evidence that they preferentially form homomers.

**Figure 1:**
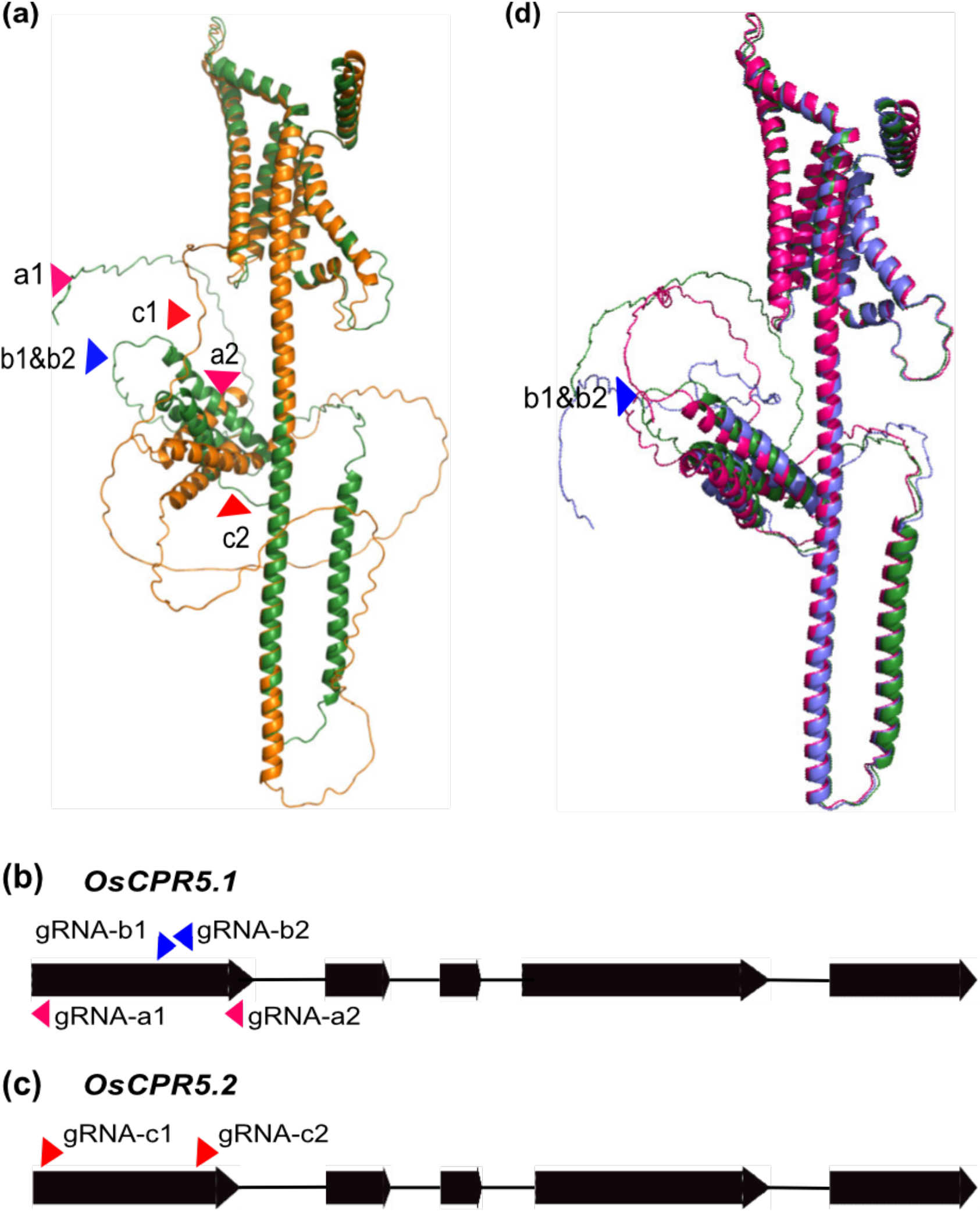
Schematic representation of *OsCPR5*.*1* and *OsCPR5*.*2* genes and of the target sites. **(a)**. Prediction of protein structures of rice CPR5. Protein structures of OsCPR5.1 and OsCPR5.2 are represented in green and orange, respectively. **(b)** The guide RNAs (gRNAs) gRNA-a1, a2, b1 and b2 were designed to target the first exon of *OsCPR5*.*1* and to induce mutations in the coding sequence. **(c)** The guide RNAs (gRNAs) gRNA-c1 and c2 were designed to target the first exon of *OsCPR5*.*2* and to induce mutations in the coding sequence. Target editing regions are indicated by pink and blue (OsCPR5.1) or red (OsCPR5.2) triangle. (**d)** Prediction of protein structures of rice OsCPR5 and in frame deletion mutants (**green:** wildtype OsCPR5.1, **hot pink:** OsCPR5.1-B8 and **blue:** OsCPR5.1-B9). CPR5 dimer predictions are shown in **Fig. S2**.

### Generation of *cpr5*.*1* mutants by genome editing

In Africa *O. glaberrima*, which is the source of most RYMV resistance genes, has been increasingly replaced by higher yielding *O. sativa* varieties that do not have the full spectrum of resistance (R) genes in their gene pool (Linares, 2002). Introgression of suitable R genes for RYMV from *O. glaberrima* into *O. sativa* remains challenging due to crossing barriers (Garavito *et al*., 2010). It may therefore be valuable to test whether loss-of-function mutations in *OsCPR5*.*1* in a *japonica* variety would confer resistance. Genome editing with CRISPR/Cas9 can efficiently be used for targeted induction of mutations, e.g., loss of function as a consequence of frameshift mutations. Since *CPR5*.*1*-based recessive RYMV resistance is associated with naturally occurring loss of function mutations, *OsCPR5*.*1* was chosen as a target for guide RNAs. To generate loss of function mutations in *OsCPR5*.*1* through CRISPR/Cas9 editing, gRNAs targeting the first exon of *OsCPR5*.*1* were designed. p-OsCPR5.1A carries gRNA-a1 and -a2, which target two sequences located 35 and 267 bp downstream the ATG to create out-of-frame deletions that would generate premature stop codons. p-OsCPR5.1B was designed to express gRNA-b1 and -b2, which target two overlapping sequences 175 bp downstream of the ATG (based on CRISPR efficiency scores and protospacer motif (PAM) sequences; Fig. 1b, c; Fig. S3, 4; Table S1). The RYMV-susceptible *O. sativa L. ssp. japonica* cv. Kitaake variety was transformed and T0 events were selected for hygromycin resistance and tested by PCR amplification to check for the presence of T-DNA. Positive T0 transformants were further subjected to sequencing for confirmation of mutations in *OsCPR5*.*1* with gene specific primers (Table S2). Sequence results were found to have bi-allelic mutations in transformants, demonstrating high efficiency of the editing process. About 50% of the mutations carried single nucleotide insertions or deletions, yet larger deletions were also observed, including a deletion of 232 nucleotides between gRNA-a1 and gRNA-a2. A subset of 13 transformants with frameshift or in frame mutations, in homo- or heterozygous states, were selfed and used for further phenotypic characterization. Progenies from 15 lines with homozygous *oscpr5*.*1* mutations and an absence of Cas9 and the *hygromycin resistance* gene, as determined by PCR, were obtained (Fig. 2a; Table S3). Ten lines (*cpr5*.*1-A1, A2, A4, B1, B2, B3, B4, B5, B6* and *B7*) carried mutations that cause frameshifts and/or early stop codons (Data S1). Three lines (*cpr5*.*1-B8, B10* and *B12*) had mutations that lead to amino-acid substitutions, and two lines (*cpr5*.*1-B9* and *cpr5*.*1-B11*) carried 42 bp in-frame deletions in the N-terminus in a domain that was predicted to be unstructured, but an otherwise intact protein (Fig. 1d; Data S1). The predicted structure of mutated OsCPR5.1 was compared to the wildtype protein structure based on OsCPR5.1 sequence in two edited lines, *cpr5*.*1-B8* and *cpr5*.*1-B9*, chosen to represent two different in frame mutations, a stretch of amino-acid change and a deletion, respectively. The *cpr5*.*1-B8* line is characterized by a substitution of eight sequential amino-acid and *cpr5*.*1-B9* by a 14-amino acids deletion. The predicted structures of the CPR5.1 variants from *cpr5*.*1-B8* and *cpr5*.*1-B9* were similar to the wildtype (Fig. 1d).

**Figure 2.**
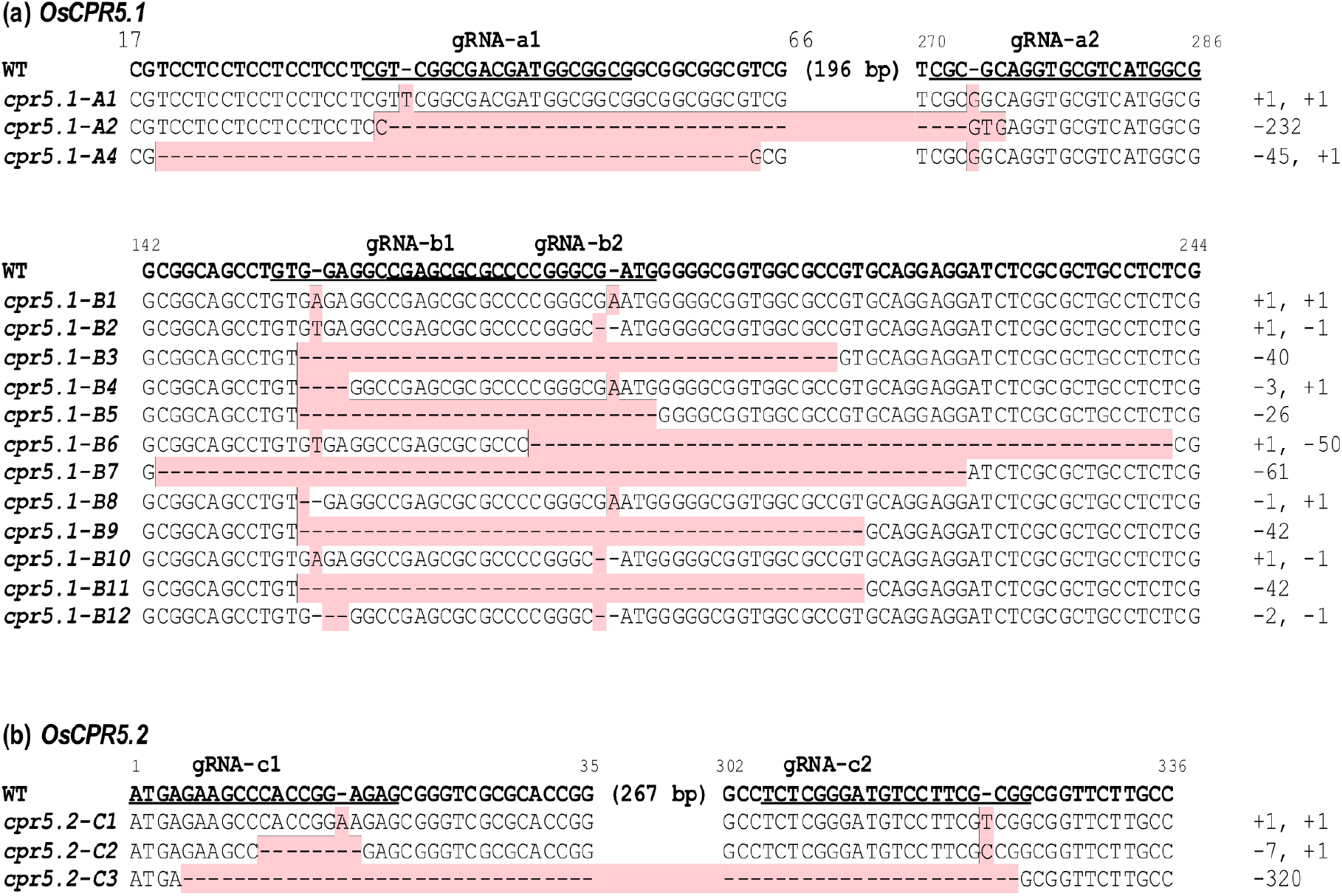
CRISPR/Cas9 mediated deletions/insertion in *OsCPR5*.*1* and *OsCPR5*.*2* in transgenic lines. gRNAs are underlined in the wildtype sequence. (**a**) Homozygous mutations in *OsCPR5*.*1*, (**b**) Homozygous mutations in *OsCPR5*.*2*. Mutations are indicated in red; “-” indicates a one-base deletion. Numbering start from ATG. The number of nucleotides inserted or deleted are indicated on the right.

### RYMV resistance of *OsCPR5*.*1* edited lines

Ten homozygous lines carrying either frameshift mutations or premature stop codons, and five lines with in-frame mutations were evaluated for RYMV resistance. Plants were mechanically inoculated with RYMV (BF1 isolate) and disease symptoms were recorded (Table S4). Wildtype Kitaake and IR64 served as susceptible controls, and two independent *oscpr5*.*1* T-DNA insertions lines as resistant controls (Pidon *et al*., 2020). Three independent experiments were performed on 3 to 17 plants per line, and for a specific line, responses were comparable (Fig. 3; Fig S5-6). Symptom observations were recorded, and virus load was determined using double antibody sandwich ELISA assays in two independent experiments. ELISA tests agreed with symptom observations two weeks after inoculation (Table S4, 5). The virus was not detected by ELISA in 4 of the 153 plants, noted as symptomatic, suggesting that in these plants, mottling and yellowing was possibly not due to the disease. Conversely, virus was detected in 16 out of 276 symptomless plants.

**Figure 3.**
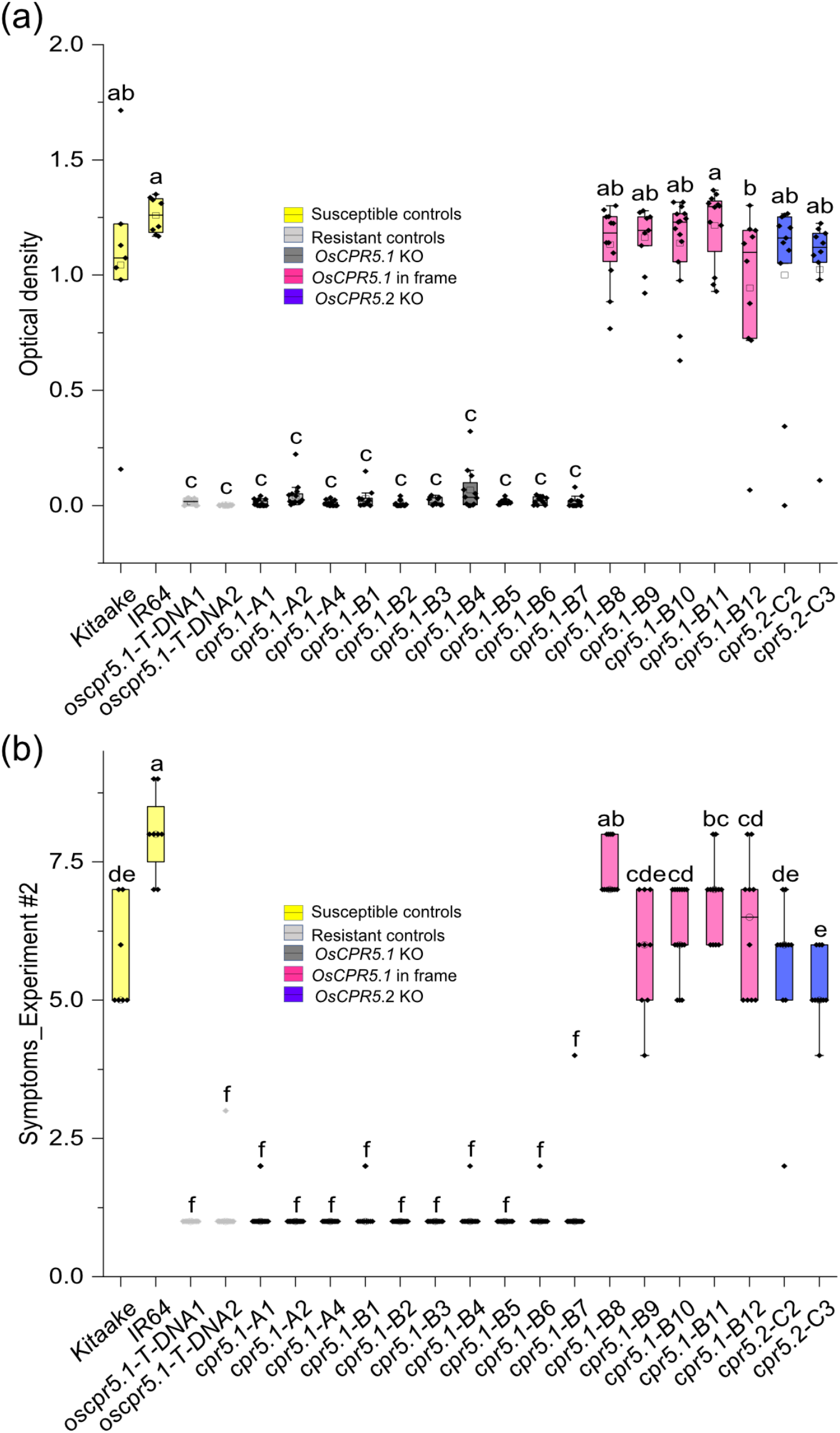
RYMV resistance evaluation in *oscpr5*.*1* and *oscpr5*.*2* mutant lines after mechanical inoculation of BF1 isolate of RYMV. Four independent experiments were performed in green house conditions and only results of experiment 2 are reported here. **(a)** Virus load in plants was detected using ELISA on leaf samples collected two weeks after inoculation. **(b)** Symptoms two weeks after inoculation were recorded using a disease scale ranging from 1 to 9. Boxes extend from 25th to 75th percentiles and display median values as center lines. Whiskers plot minimum and maximum values, asterisks indicate individual data points and detected by one-way ANOVA followed by Tukey’s test, different letters indicate significant differences (p< 0.05).

Lines with frameshift mutation or premature stop codons showed full resistance. Only 8 out of 564 plants showed mottling and chlorosis two weeks after inoculation (Fig. 3a, b; Fig S5, 6) Loss-of-function of *OsCPR5*.*1* is thus sufficient to confer RYMV resistance in a *japonica* variety (Fig. 3a, b). On the contrary, symptoms were observed in 265 out of 267 plants with in frame mutations in *OsCPR5*.*1* indicating that lines with in frame mutations were susceptible (Fig. 1d). Most likely, modification of the possibly unstructured domain at the N-terminus is not relevant for virulence of RYMV (Fig. 1d). Interestingly, *oscpr5*.*1-B8* and *oscpr5*.*1-B10* showed more severe symptoms compared to Kitaake wild-type in all the four experiments, while comparable symptom development was observed in two experiments for *oscpr5*.*1-B9, - B11* and *-B12*. This result may indicate a hyper-susceptibility associated with the in-frame deletions. Taken together, loss-of-function of *OsCPR5*.*1* is sufficient to confer RYMV resistance to a *O. sativa ssp. japonica* variety.

### Sheath blight disease resistance evaluation

Seven homozygous frameshift mutants, one in frame mutant of *oscpr5*.*1* and Kitaake wildtype were evaluated for sheath blight disease resistance. Plants were inoculated with the virulent *R. solani* strain RSY-04 and observations were recorded. Based on lesion length produced on rice leaves, all tested *oscpr5*.*1* mutants were susceptible, and lesions were similar as in Kitaake wildtype (Fig. S7).

### Agro-morphological characters of *oscpr5*.*1* mutants

Arabidopsis *atcpr5* mutants showed increased resistance against diverse pathogens viz., *Pseudomonas syringae pv Pst DC3000, Pseudomonas syringae pv maculicola ES4326, Erisyphe cruciferarum*, and *Hyaloperonospora arabidopsidis*, as well as compromised growth, indicating that *AtCPR5* plays important roles in broad spectrum pathogen susceptibility and in the physiology of uninfected plants (Höwing *et al*., 2017; Wang *et al*., 2017). To test whether *oscpr5*.*1*-mediated RYMV resistance is associated to a reduction of plant growth, we evaluated key agro-morphological characters of the mutants in the absence of the virus in three independent experiments at HHU (Table S6). Plant height, panicle length, reproductive tillers/plant and grain weight were evaluated in seven frameshift mutants and five in-frame mutants and no significant differences from wildtype Kitaake were found (Fig. 4; Fig. S8, 9). Only *oscpr5*.*1-B4* showed defects, likely due to second site mutations that may impact panicle length. Three frameshift mutant lines *oscpr5*.*1-B5, oscpr5*.*1-B6* and *oscpr5*.*1-B7* showed increased reproductive tiller/plant and grain weight/plant (Fig. 4). In summary, *oscpr5*.*1* edited frameshift and in frame mutants were generally not significantly different from Kitaake wildtype regarding various growth parameters.

**Figure 4.**
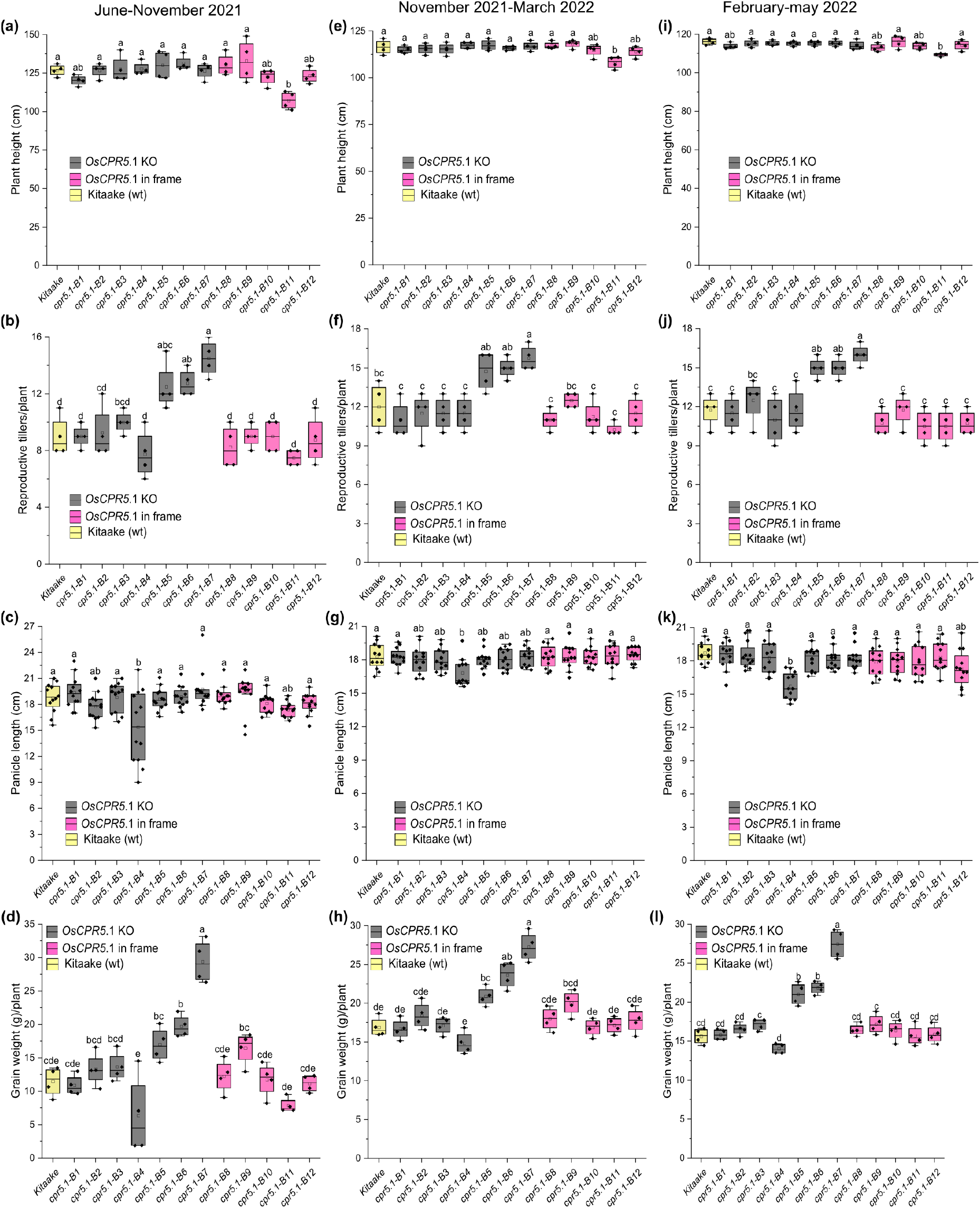
Agro-morphological characters of *OsCPR5*.*1* edited mutants. Across three independent experiments, no reduction in plant height (cm), reproductive tiller number/plant and grain weight (g)/plant were observed between the wildtype control Kitaake and the *knockout* mutant lines *oscpr5*.*1-B1, 2, 3, 5, 6*, and *7*. B5-B7 even showed increased plant height and grain weight. Line *oscpr5*.*1-B4* likely contains second site mutations that may impact all three phenotypes. Boxes extend from 25th to 75th percentiles and display median values as center lines. Whiskers plot minimum and maximum values, asterisks indicate individual data points and detected by one-way ANOVA followed by Tukey’s test, different letters indicate significant differences (p< 0.05). Data were generated from three independent experiments conducted in HHU greenhouses under controlled conditions.

### The role of *OsCPR5*.*2* in RYMV resistance

OsCPR5.1 and OsCPR5.2 share 56% of identity and were predicted to be structurally highly similar (Fig. 1c; Fig. S10). Although natural *oscpr5*.*2* alleles of have not been described in the context of RYMV resistance, it is conceivable that *oscpr5*.*2* knockout mutants could also confer RYMV resistance. To explore the role of *OsCPR5*.*2*, in particular for RYMV resistance, two sgRNAs were designed to target sequences in the first exon of *OsCPR5*.*2* (Fig. 1c; Table S1). Seventeen independent T0 transformants of the *O. sativa L. ssp. japonica cv*. Kitaake variety were selected based on hygromycin resistance and PCR amplification, and then mutations in *OsCPR5*.*2* were confirmed by sequencing using gene specific primers (Table S2). Most lines carried bi-allelic mutations in *OsCPR5*.*2*. Three transformants with frameshift mutations or premature stop codons, leading to loss-of-function alleles, were advanced to T2 homozygous progenies (Fig. 2b; Table S7). Three lines with frameshift mutations (*oscpr5*.*2-C1, oscpr5*.*2-C2* and *oscpr5*.*2-C3*) were evaluated for RYMV resistance (Data S2). All three mutants were susceptible to RYMV and not significantly different from the wildtype Kitaake control regarding symptoms and virus load (Fig. 3a-b; Fig. S6, 7). The agro-morphological characters of *OsCPR5*.*2* frameshift mutants were analyzed, as described above for *OsCPR5*.*1* frameshift mutant lines (Fig. S11; Table S8). Plant height and grain weight of *oscpr5*.*2-C3* were reduced compared to the Kitaake wildtype, but *oscpr5*.*2-C1* and *oscpr5*.*2-C2* mutants were not significantly different to Kitaake wildtype for any of the three traits evaluated. The results obtained of *oscpr5*.*2-C1* and *oscpr5*.*2-C2* suggest that loss of function of OsCPR5.2 did not affect growth (Fig. S11), and defects observed in *oscpr5*.*2-C3* were likely due to second site mutations.

## Discussion

To evaluate whether genome editing of the two rice *CPR5* homologs can be used to confer resistance to RYMV, a major biotic constraint affecting rice production in sSA (Odongo *et al*., 2021), two nucleoporin paralogs were edited in the japonica rice variety Kitaake (Fig. S12). While homozygous loss of function alleles carrying premature stop codons in *OsCPR5*.*1* led to full resistance, loss of function mutations in *OsCPR5*.*2* did not show a detectable increase in resistance. Interestingly, deletions in the N-terminal part of *OsCPR5*.*1* appeared to trigger even increased susceptibility. Thus, despite the predicted structural similarity and the ability of the two paralogs to complement the mutant phenotype of the Arabidopsis *atcpr5* mutant (Mei *et al*., 2022), *OsCPR5*.*1* and *OsCPR5*.*2* have distinct functions with respect to susceptibility to RYMV (Fig.3; Table 1). Notably, neither *oscpr5*.*1*, nor *oscpr5*.*2* single *knockout* mutants showed apparent penalties under greenhouse conditions. Also, naturally occurring *rymv2* variants did not show penalties, thus *OsCPR5*.*2* may compensate for loss of the basic nucleoporin functions of OsCPR5.1. It will be interesting to analyze *oscpr5*.*1 oscpr5*.*2* double mutants to evaluate whether the combination of mutations lead to growth defects similar to those observed in the Arabidopsis *atcpr5* mutant. Taken together, our data indicate that genome editing events that cause loss of function in *OsCPR5*.*1* provide an effective approach to introduce resistance to RYMV into elite varieties of rice, which otherwise cannot be introduced easily by conventional breeding or marker assisted selection due to sterility barriers between *O. glaberrima* and *O. sativa* L. (Garavito *et al*., 2010). Since some RYMV isolates are able to overcome individual R-genes against RYMV in experimental conditions (not observed in the field), editing may provide an effective tool to stack multiple R genes and obtain more robust resistance (Bonnamy *et al*., 2023; Hébrard *et al*., 2018; Pinel-Galzi *et al*., 2016). Other groups recently showed that genome editing can be used to generate resistance against other viruses by targeting translation initiation factors, known as major susceptibility factors in plant/virus interactions (Atarashi *et al*., 2020; Kuroiwa *et al*., 2022; Macovei *et al*., 2018; Pechar *et al*., 2022).

**Table 1.** Summary of features of *oscpr5*.*1* and *oscpr5*.*2* edited lines.

| Genotype | Mutation | Symptoms | ELISA | Plant height % | Tillers/plant % | Panicle length % | Grain weight % |
| --- | --- | --- | --- | --- | --- | --- | --- |
| Kitaake | Wildtype | + | S | 100 | 100 | 100 | 100 |
| IR64 | Wildtype | + | S | NA | NA | NA | NA |
| *oscpr5.1-A1* | Stop codon after aa 33 | - | R | NA | NA | NA | NA |
| *oscpr5.1-A2* | Stop codon after aa 15 | - | R | NA | NA | NA | NA |
| *oscpr5.1-A4* | Stop codon after aa 90 | - | R | NA | NA | NA | NA |
| *oscpr5.1-B1* | Stop codon after aa 93 | - | R | 96.9 | 96.2 | 97.2 | 98.8 |
| *oscpr5.1-B2* | Stop codon after aa 59 | - | R | 98.0 | 101.5 | 96.0 | 109.7 |
| *oscpr5.1-B3* | Stop codon after aa 79 | - | R | 100.5 | 103.1 | 93.3 | 108.8 |
| *oscpr5.1-B4* | Stop codon after aa 104 | - | R | 98.8 | 93.1 | 90.2 | 80.1 |
| *oscpr5.1-B5* | Stop codon after aa 96 | - | R | 101.0 | 122.9 | 95.2 | 133.8 |
| *oscpr5.1-B6* | Stop codon after aa 59 | - | R | 100.2 | 129.8 | 96.0 | 148.2 |
| *oscpr5.1-B7* | Stop codon after aa 72 | - | R | 98.8 | 138.2 | 103.8 | 191.1 |
| *oscpr5.1-B8* | Substitution 60-67 aa | + | S | 99.8 | 106.9 | 95.5 | 106.1 |
| *oscpr5.1-B9* | deletion 59-73 aa | + | S | 102.2 | 96.2 | 103.5 | 122.0 |
| *oscpr5.1-B10* | Substitution 60-67 aa | + | S | 95.7 | 96.2 | 92.9 | 102.2 |
| *oscpr5.1-B11* | deletion 59-73 aa | + | S | 92.2 | 88.5 | 98.3 | 92.6 |
| *oscpr5.1-B12* | Substitution 60-67 aa | + | S | 96.3 | 93.9 | 94.8 | 101.8 |
| Kitaake (IRD) | wildtype | + | S | 100 | 100 | NA | 100 |
| *oscpr5.2-C1* (IRD) | Stop codon after 138 aa | + | S | 98.2 | 101.0 | NA | 101.1 |
| *oscpr5.2-C2* (IRD) | Stop codon after 19 aa | + | S | 99.7 | 96.2 | NA | 102.7 |
| *oscpr5.2-C3* (IRD) | Stop codon after 8 aa | + | S | 91.3 | 93.5 | NA | 80.4 |

Values are from average of all experiments; +: positive, -: negative, R: resistant (blue), S: susceptible, NA: not analyzed. Yellow and green indicate significant deviation from controls.

### Potential role of *OsCPR5*.*1* in RYMV susceptibility

Since loss of the function leads to resistance, *OsCPR5*.*1* enables the virus to become virulent. Thus, *OsCPR5*.*1* serves as a recessive susceptibility gene. All CPR5 proteins appear to encode integral membrane proteins with five predicted transmembrane helices and a large soluble domain and likely function as membrane-anchored nucleoporins. AtCPR5 had been shown to interact genetically and physically with at least three other nucleoporins (Nup85, Nup96, NUP160) as well as the karyopherin Exportin-4 (Gu *et al*., 2016; Xu *et al*., 2021). AtCPR5 can form homodimers, which are disrupted when effector-triggered immunity (ETI) is activated, resulting in conformational changes that trigger the release of cyclin-dependent kinase inhibitors and other factors involved in defense ultimately leading to programmed cell death and hypersensitive responses as well as other defenses that block the reproduction/replication and limit the spread of the pathogen (Xu *et al*., 2021). This fundamental pathway therefore is relevant to broad spectrum resistance/susceptibility to bacterial, fungal and viral diseases. Besides, since CPR5, at least in Arabidopsis, appears to be a component of the nuclear pore, one may speculate that the nucleus is required for effective virus replication. One may propose two alternative hypotheses:

i. Resistance against RYMV in *OsCPR5*.*1* loss of function mutants could possibly result from the constitutive activation of defense, including the salicylic acid (SA) signaling pathway (Bowling *et al*., 1997). Consistent with this hypothesis, *atcpr5* mutants are resistant to the bacterial pathogen *P. syringae* and the oomycete *H. arabidopsidis* (Bowling *et al*., 1997). CPR5 was found to bind to the N-terminus of the ethylene receptor ETR1, and is thought to regulate the export of ethylene-related mRNAs from the nucleus (Chen *et al*., 2022). In the case of virus, SA is known to induce effective resistance to a wide range of plant virus by inhibiting viral replication, blocking intercellular transport and systemic spread (Chivasa *et al*., 1997; Métraux *et al*., 1990; Murphy *et al*., 2020; Yalpani *et al*., 1993). For instance, cauliflower mosaic virus failed to systemically spread in *atcpr5* mutants (Love *et al*., 2007). It is generally assumed that constitutive defense activation brings trade-offs; consistent with the massive penalty observed in *atcpr5* mutants in Arabidopsis. Since in *O. glaberrima* and *O. sativa oscpr5*.*1* mutants do not have apparent penalties, it is rather unlikely that the *rymv2* resistance is due to defense priming. Moreover, *rymv2* would be expected to confer broad disease resistance, which was however not been observed in preliminary tests using *Rhizoctonia solani* (sheath blight) *Magnaporthe grisea* (blast), and *Xanthomonas oryzae* pv. *oryzae* (bacterial leaf blight) (Fig. S7 and unpublished results).
ii. Alternatively, CPR5.1 could play a specific role that is required for RYMV virulence. RNA virus replication does not strictly depend on import into the nucleus, but various viruses, including RNA viruses, require proteins from the nucleus and disrupt the nucleocytoplasmic transport or target the nucleolus to promote their own replication (Flather and Semler, 2015; Walker and Ghildyal, 2017). Opalka et al., (1998) showed that RYMV virions can be detected in the nuclei of mesophyll cells (Opalka *et al*., 1998). However, at present, it is not known whether RYMV RNA enters the nucleus and whether virus replication is impaired in the *cpr5*.*1* mutants.

### No obvious role of *OsCPR5*.*2* in RYMV susceptibility

By contrast to *oscpr5*.*1* loss of function mutants, *oscpr5*.*2* mutants did not show detectable RYMV resistance, implicating that *OsCPR5*.*2* is not directly involved in the interaction with the virus, at least when *OsCPR5*.*1* is functional. However, the functional redundancy between paralogs of virus susceptibility genes can affect the resistance durability. Resistance-breakdown can result from the restoration of the interaction of the virus with the susceptibility factor when resistance alleles are characterized by substitutions or small deletions (Charron *et al*., 2008; Hébrard *et al*., 2010), which is not possible when resistance is conferred by knockout mutations. In that case, some viruses can recruit paralogs or isoforms of the missing susceptibility factor, as documented in the potyvirus/eIF4E and eIF(iso)4E interactions (Bastet *et al*., 2018; Takakura *et al*., 2018). Pinel-Galzi *et al*. (2016) reported that some RYMV isolates can overcome *RYMV2*-mediated resistance by acquiring mutations in the membrane anchor domain of the P2a polyprotein (Pinel-Galzi *et al*., 2016). Those mutations could possibly result from an adaptation of the virus to *OsCPR5*.*2* when *OsCPR5*.*1* is defective. Besides, *OsCPR5*.*1* and *OsCPR5*.*2* genes are broadly expressed in all plant tissues (Fig. S1); one thus may hypothesize that they can form heterodimers. However, the heterodimerization, if it occurs, does not appear to be relevant to susceptibility, since *oscpr5*.*2* mutants do not show a substantial increase in resistance. To dissect the resistance mechanism, it will be important to determine the subcellular localization of the two isoforms, the ability to dimerize, the effect of the virus on localization and dimerization as well as the effect of loss of CPR5.1 function on viral RNA entry into the nucleus and viral replication.

### Summary and outlook

Our results confirmed that the disruption of *OsCPR5*.*1* in the *O. sativa L. ssp. japonica cv*. Kitaake can confer high level of resistance to RYMV isolate BF1 without any yield penalty, at least under greenhouse conditions. Novel mutations can effectively be introduced by CRISPR/Cas9 into rice cultivars to obtain resistance against the RYMV and might help in accelerated breeding applications to develop resistance in popular cultivars in sSA. The next step will be to transfer this approach to an elite variety better adapted to rice culture in rainfed lowland and irrigated areas of Africa, where rice production is particularly affected by RYMV disease. Suitable transformation protocols have been established (Luu *et al*., 2020). To obtain more robust resistance, it will be important to stack the three known resistance genes to be able to also protect against new RYMV variants that could break the resistance genes if deployed individually. The resistance-breaking of *RYMV2* is associated to mutations in the membrane anchor domain of the P2a polyproteine (Pinel-Galzi et al, 2016). *RYMV1* encodes the translation initiation factor eIF(iso)4G1 that is recruited by the virus and directly interacts with the viral Protein genome-linked (VPg) covalently fused to the 5′ ends of the positive strand viral RNA (Hébrard *et al*., 2010) (Albar *et al*., 2006). In incompatible interactions, non-synonymous mutations in eIF(iso)4G1 impair the interaction with the VPg causing a resistant phenotype and mutations in the VPg allow virus adaptation and resistance-breaking (Hébrard *et al*., 2006, 2010). *RYMV3* is a classical NB-LRR that recognizes the coat protein of the virus and likely acts upstream of CPR5.1 (Bonnamy *et al*., 2023). Independent viral determinants are thus involved in the resistance-breaking of the different resistance genes. stacking of suitable alleles of all three S/R genes will likely help to increase robustness. Further clarification of the susceptibility and resistance mechanisms might help to design robust resistance. Subsequently, field trials will be required to validate that resistance is maintained under the local conditions in Africa, and to evaluate possible penalties in the relevant agro-ecosystems.

## Methods

### Plasmid construct and generation of transgenic rice lines

Mutations in the target genes *OsCPR5*.*1* (*LOC_Os01g68970*.*1* gene model on the reference Nipponbare sequence) and *OsCPR5*.*2* (*LOC_Os02g53070*.*1*) were generated through CRISPR/Cas9 editing in *Oryza sativa* cv. Kitaake, which is susceptible to RYMV. Two pairs of gRNAs (gRNA-a1/gRNA-a2 and gRNA-b1/gRNA-b2) were designed on the first exon of *OsCPR5*.*1* and one pair of gRNAs (gRNA-c1/gRNA-c2) on the first exon of *OsCPR5*.*2* gene using online tools (https://cctop.cos.uni-heidelberg.de/; https://www.benchling.com/; Fig. 1a-b; Table S1).

For the gRNA-b1/gRNA-b2 pair, single-stranded oligonucleotides were synthesized with adapter sequences for forward (5’-GCAG-3’) and reverse primers (5’-AAAC-3’) (IDT). The single-stranded DNA oligos were annealed to produce double-stranded oligonucleotides. Two double-stranded oligonucleotides were individually inserted into the *BsmBI*-digested pTLNt-gRNA-1 to T2 to generate two tRNA-gRNA units. Two tRNA–gRNA units were transferred into *BsmBI*-digested pENTR4-U6.1PccdB using the Golden Gate ligation method. The cassettes containing two tRNA-gRNA units under control of U6.1P were finally mobilized to a binary vector pBY02-ZmUbiP:SpCas9 (Char *et al*., 2017; Zhou *et al*., 2014) using Gateway LR Clonase (Thermo Fisher Scientific) and designated as p-OsCPR5.1-B (Fig. S4).

For the two other gRNA pairs, two cassettes containing the two gRNAs under the control of the rice promoters U6.1p and U6.2p, the sgRNA1 and attL1 and attL2 motives, were synthesized. These cassettes were sub-cloned into a modified binary cas9 vector pH-Ubi-cas9-7 (Fayos *et al*., 2020; Miao *et al*., 2013) in a Gateway LR reaction (Invitrogen) to generate p-OsCPR5.1-A and p-OsCPR5.1-C constructs (Fig. S3; Fig. S10).

*Agrobacterium tumefaciens* strain EHA105 was transformed by electroporation with Cas9/gRNA-expressing binary vectors. Rice transformation was performed by *A. tumefaciens* as previously described (Blanvillain-Baufumé *et al*., 2017; Sallaud *et al*., 2003; Wang *et al*., 2017). Transgenic seedlings at 5-leaf stage were transferred to greenhouse.

### Genotyping and sequencing of edited lines

Genomic DNA was extracted using the peqGOLD Plant DNA Mini Kit (PeqGold, VWR International GmbH, Darmstadt, Germany) for lines transformed with pOsCPR5.1-B construct while extraction was done according to (Edwards *et al*., 1991) for lines transformed with p-OsCPR5.1-A and p-OsCPR5.1-C constructs. Presence of T-DNA insertion was detected by specific amplification of SpCas9 and hygromycin resistance gene fragments. Primer pairs Os-Cas9-F/OsCas9-R and Hygro-IIF/Hygro-IIR were used for lines transformed with pOsCPR5.1-B construct, and primer pairs Cas9-F(IRD)/cas9-R(IRD) and Hygro-F(IRD)/Hygro-R(IRD) for lines transformed with p-OsCPR5.1-A and p-OsCPR5.1-C constructs (Table S2). Amplifications were performed using the High-Fidelity Phusion PCR Master Mix (NEB, Frankfurt am Main, Germany) or the GoTaq G2 polymerase (Promega, Madison, US) following the manufacturer’s instructions. Genome edition was analyzed by sequencing of amplification products containing the target loci. Lines obtained with constructs p-OsCPR5.1-A, p-OsCPR5.1-B, p-OsCPR5.1-C were amplified with gene specific primers pairs (Table S2), respectively. Sequencing was subcontracted to Genewiz or Microsynth. Superimposed sequencing chromatograms were analyzed with DSDecodeM to decode heterozygous profiles and validated manually.

### RYMV resistance evaluation

Rice plants were grown at IRD in plastic trays in greenhouse, at a 28°C/22°C day/night temperature, 70% humidity and a 14h light/24h photoperiod. The wildtype *O. sativa L. ssp. japonica* cv. Kitaake and *O. sativa indica* IR64 were used as susceptible controls; T-DNA insertion lines in *O. sativa L. ssp. japonica* cv. Hwayoung and Dongjin (Pidon *et al*., 2020) were used as resistant controls. Four independent experiments were performed. Plants were mechanically inoculated with RYMV about 2 weeks after sowing as described in (Pinel-Galzi *et al*., 2018) with the BF1 isolate of RYMV. Resistance reaction was assessed based on the presence/absence of symptoms, completed with double-antibody sandwich (DAS-) ELISA for experiments 1 and 2. Symptoms were observed on the leaves emerged after inoculation 2 weeks after inoculation. Samples from the most recently emerged leaf were collected one or two weeks after inoculation for experiments 2 and 1, respectively, and virus load was estimated by direct double-antibody sandwich (DAS-) ELISA, as described by (Pinel-Galzi *et al*., 2018), using a polyclonal antiserum directed against an RYMV isolate from Madagascar (N’Guessan *et al*., 2000).

### Sheath blight disease resistance evaluation

To explore resistance of *oscpr5*.*1* mutants to *R. solani*, seven *oscpr5*.*1* frameshift mutants (*oscpr5*.*1, B1-B7*), one in frame mutant (*oscpr5*.*1-B11*) and wildtype Kitaake were grown at HHU. 30-day-old plants were used for inoculation with *R. solani* (RSY-04). Individual sclerotia from three-day old fresh *R. solani* grown on Potato Dextrose Agar (PDA) were inoculated onto individual rice leaves and incubated at 27 °C. Phenotypic observations (8 plants/mutant) were recorded 5 days post inoculation. Notably, this experiment was performed only once (Fig. S7).

### CPR5 structure predictions

Protein structures for AtCPR5 (UniProt ID: Q9LV85), OsCPR5.1 (UniProt ID: Q5JLB2), OsCPR5.2 (UniProt ID: Q6ZH55) selected mutants and possible dimers were generated using the AlphaFold Colab notebook interface (https://colab.research.google.com/github/deepmind/alphafold/blob/main/notebooks/AlphaFol d.ipynb) based on the AlphaFold v2 structure prediction program (Jumper *et al*., 2021; Mirdita *et al*., 2022) using a selected portion of the BFD database (https://bfd.mmseqs.com). Accuracy in predicted structures might therefore differ slightly from those obtained from the full AlphaFold system. We note that these are predictions that may not reflect the actual *in vivo* structures.

### Plant growth and analysis of agro-morphological characters

Plants obtained with pOsCPR5.1-B construct were grown at HHU in greenhouse conditions at 30 °C during the day time and 25 °C during the night time with 60-70% relative humidity. Light conditions in the greenhouse are determined by natural daylight and additional lamplight (8/16 day/night photoperiod). Plants obtained with pOsCPR5.1-A and -C constructs were grown at IRD in greenhouse conditions with a 28°C/22°C day/night temperature, 70% humidity and a 14h light/24h photoperiod. To determine the agro-morphology of *oscpr5* lines, four to five plants of each line were grown in greenhouses and measured for different morphological characters viz., plant height, number of reproductive tillers, panicle length and grain weight. Three independent experiments were performed for each set of lines.

## Acknowledgements

We would like to thank Prof. Bing Yang for providing CRISPR system. We would like to thank Perlina Lim for excellent technical assistance. We are grateful to Emilie Thomas and Harold Chrestin for their help during greenhouses experiments, and to Agnes Pinel-Galzi for providing anti-RYMV antibodies. The master student Oscar Main contributed to preliminary phenotypic experiments. This work was supported by grants from the Bill and Melinda Gates Foundation to HHU, a Marie Skłodowska-Curie fellowship under the European Union’s Horizon 2020 research and innovation programme (“PDgate” No. 101023981) to MM, Deutsche Forschungsgemeinschaft (DFG, German Research Foundation) under Germany’s Excellence Strategy – EXC-2048/1 – project ID 390686111 (CEPLAS), and an Alexander von Humboldt Professorship (WF). YA was supported by fellowships from the Alexander von Humboldt Foundation and CEPLAS. At IRD, this work was financially supported by the CGIAR Research Program on rice agri-food systems (RICE, 2017–2022).

## Conflict of interest

The authors declare no conflict of interest.

## Authors contribution

YA, SC, WBF and LA developed the concept. YA, FA, MS, SC, and LA performed experiments. YA, FA, LA, MMW and VSL have performed in silico analyses. YA, LA, MM and WBF, have written the manuscript. All authors have given approval to the final version of the manuscript.

## Supporting information

### Supporting Data

**Data S1**. Predicted amino acid sequence of *Os*CPR5.1 in wildtype and in mutants.

**Data S2**. Predicted amino acid sequence of *Os*CPR5.2 in wildtype and mutants.

### Supporting Figures

**Figure S1**. Tissue specific and developmental stages expression levels of *OsCPR5*.*1* and *OsCPR5*.*2*.

**Figure S2**. Alphafold prediction of AtCPR5, *Os*CPR5.1 and *Os*CPR5.2 dimer conformations.

**Figure S3**. Map of the binary CRISPR/Cas9 vector p-OsCPR5.1-A.

**Figure S4**. Map of the binary CRISPR/Cas9 vector p-OsCPR5.1-B.

**Figure S5**. Disease resistance phenotypic reaction of *oscpr5*.*1* and *oscpr5*.*2* mutants.

**Figure S6**. Symptoms of *oscpr5*.*1* and *oscpr5*.*2* mutant lines two weeks after inoculation with BF1 isolate of RYMV.

**Figure S7**. Disease phenotypic reaction of *oscpr5*.*1* mutants with *R. solani* AG1-1A.

**Figure S8**. Phenotypic characters of *oscpr5*.*1* mutants.

**Figure S9**. Panicle characters of *oscpr5*.*1* mutant plants.

**Figure S10**. Map of the binary CRISPR/Cas9 vector p-OsCPR5.2-C.

**Figure S11**. Morphological characters of *oscpr5*.*2* frameshift mutant plants.

**Figure S12**. Alignment of CPR5 homologs compared to *Arabidopsis thaliana*.

### Supporting Tables

**Table S1**. List of gRNAs used to develop CRISPR/Cas9 mediated mutations of *OsCPR5*.*1* and *OsCPR5*.*2* in Kitaake

**Table S2**. List of primers used in the present study

**Table S3:** List of CRISPR/Cas9-induced insertions or deletions in T2 homozygous plants of *OsCPR5*.*1*

**Table S4**. RYMV disease resistance phenotypic reaction of *oscpr5*.*1* and *oscpr5*.*2* (**additional file**).

**Table S5:** Virus detection using ELISA in symptomatic and symptomless plants at two weeks after inoculation with BF1 isolate of RYMV

**Table S6**. Agro-morphological characters of *oscpr5*.*1* (**additional file**).

**Table S7:** List of CRISPR/Cas9-induced deletions in T2 homozygous plants of *oscpr5*.*2*.

**Table S8**. Agro-morphological characters of *oscpr5*.*2* (**additional file**).

## Notes

### Competing Interest Statement

The authors have declared no competing interest.

