## Supplementary Data for "*Rice Yellow Mottle Virus* resistance by genome editing of the *Oryza sativa* L. ssp. japonica nucleoporin gene *OsCPR5.1* but not *OsCPR5.2*"

* Equal contribution

**List of Content**

**Supporting Data**

**Data S1.** Predicted amino acid sequence of *Os*CPR5.1 in wildtype and in mutants.

**Data S2**. Predicted amino acid sequence of *Os*CPR5.2 in wildtype and mutants.

**Supporting Figures**

**Figure S1.** Tissue specific and developmental stages expression levels of *OsCPR5.1* and *OsCPR5.2*.

**Figure S2.** Alphafold prediction of AtCPR5, *Os*CPR5.1 and *Os*CPR5.2 dimer conformations.

**Figure S3.** Map of the binary CRISPR/Cas9 vector p-OsCPR5.1-A.

**Figure S4.** Map of the binary CRISPR/Cas9 vector p-OsCPR5.1-B.

**Figure S5.** Disease resistance phenotypic reaction of *oscpr5.1* and *oscpr5.2* mutants.

**Figure S6**. Symptoms of *oscpr5.1* and *oscpr5.2* mutant lines two weeks after inoculation with BF1 isolate of RYMV.

**Figure S7.** Disease phenotypic reaction of *oscpr5.1* mutants with *R. solani* AG1-1A.

**Figure S8.** Phenotypic characters of *oscpr5.1* mutants.

**Figure S9.** Panicle characters of *oscpr5.1* mutant plants.

**Figure S10.** Map of the binary CRISPR/Cas9 vector p-OsCPR5.2-C.

**Figure S11.** Morphological characters of *oscpr5.2* frameshift mutant plants.

**Figure S12.** Alignment of CPR5 homologs compared to *Arabidopsis thaliana*.

**Supporting Tables**

**Table S1.** List of gRNAs used to develop CRISPR/Cas9 mediated mutations of *OsCPR5.1* and *OsCPR5.2* in Kitaake

**Table S2.** List of primers used in the present study

**Table S3:** List of CRISPR/Cas9-induced insertions or deletions in T2 homozygous plants of *OsCPR5.1*

**Table S4.** RYMV disease resistance phenotypic reaction of *oscpr5.1* and *oscpr5.2* (**additional file**).

**Table S5:** Virus detection using ELISA in symptomatic and symptomless plants at two weeks after inoculation with BF1 isolate of RYMV

**Table S6.** Agro-morphological characters of *oscpr5.1* (**additional file**).

**Table S7:** List of CRISPR/Cas9-induced deletions in T2 homozygous plants of *oscpr5.2*.

**Table S8.** Agro-morphological characters of *oscpr5.2* (**additional file**).

**Data S1.** Predicted amino acid sequence of *Os*CPR5.1 in wildtype and in mutants. Amino acid sequences similar to the wildtype *Os*CPR5.1 sequence are in black font, differences to the wildtype *Os*CPR5.1 sequence are in red.

**>*Os*CPR5.1**

MDAAAASSSSSSSATMAAAAASAAEASLSGSPASSRNARHRQRKGVRLRMLRRRGRQPVEAERAPGDG
GGGAVQEDLALPLGMSFAAVLAQVINTKNISGQRLHPDFLSKICTSAVKESLTNIYGDSSNSFIKNFE
KSFSSTFRTLHLVNEIPVNERSHIPECSFKHDDSVAVDSLSSSDLQNQTNRIEHDLVNTVESQLVLFA
SDNQQLTHLRHSRSSPEADNRILNAIDRSNELKEFEIGLTMRKLQLKQSQLALSSHSHMLEKIKLSFG
FQKASFKGEKFKTRMQETRDAEILRTLIDFLVSAVIVMSACFGYGTYIYSYQRITDVTSACSATSKGS
KSWWMPNSVSNFSSGFLFLRCHVIAVTRMCFGILMILAIAWLAFQRSSTTGSNMPITFNLILLGIICG
FAGRFCTNTLGGDGNTWLMYWEVLCSIHLLGNLFPSLLYHVLHGPISVSHREQVVWLPYWVRRCLFYA
AVGLILPALTGLLPFASLSDWKDHFVEEIKSIVIGDKIEA*

**>oscpr5.1-A1**

MDAAAASSSSSFGDDGGGGGVGRGGVVVRLACFFEERSPSAA*

**>oscpr5.1-A2**

MDAAAASSSSSVRL*IQRIFQDKDYILISSPRSVHQQSRNL*

**>oscpr5.1-A3**

MDAAAASSSSSYGDDGGGGGVGRGGVVVRLACFFEERSPSAA*

**>oscpr5.1-A4**

MDAAAAAAA...............EASLSGSPASSRNARHRQRKGVRLRMLRRRGRQPVEAERAPGDG
GGGAVQEDLALPLGMSFAAVLAAGYKYKEYFRTKITS*

**>oscpr5.1-B1**

MDAAAASSSSSSSATMAAAAASAAEASLSGSPASSRNARHRQRKGVRLRMLRRRGRQPVRGRARPGRM
GAVAPCRRISRCLSECPSPPSSRRL*

**>oscpr5.1-B2**

MDAAAASSSSSSSATMAAAAASAAEASLSGSPASSRNARHRQRKGVRLRMLRRRGRQPV*

**>oscpr5.1-B3**

MDAAAASSSSSSSATMAAAAASAAEASLSGSPASSRNARHRQRKGVRLRMLRRRGRQPVCRRISRCLS
ECPSPPSSRRL*

**>oscpr5.1-B4**

MDAAAASSSSSSSATMAAAAASAAEASLSGSPASSRNARHRQRKGVRLRMLRRRGRQPVAERAPGEWG
RWRRAGGSRAASRNVLRRRPRAGYKYKEYFRTKITS*

**>oscpr5.1-B5**

MDAAAASSSSSSSATMAAAAASAAEASLSGSPASSRNARHRQRKGVRLRMLRRRGRQPVGRWRRAGGS
RAASRNVLRRRPRAGYKYKEYFRTKITS*

**>oscpr5.1-B6**

MDAAAASSSSSSSATMAAAAASAAEASLSGSPASSRNARHRQRKGVRLRMLRRRGRQPV*

**>oscpr5.1-B7**

MDAAAASSSSSSSATMAAAAASAAEASLSGSPASSRNARHRQRKGVRRRMLRRRGISRCLSECPSPPS
SRRL*

**>oscpr5.1-B8**

MDAAAASSSSSSSATMAAAAASAAEASLSGSPASSRNARHRQRKGVRLRMLRRRGRQPVRPSAPRANG
GGGAVQEDLALPLGMSFAAVLAQVINTKNISGQRLHPDFLSKICTSAVKESLTNIYGDSSNSFIKNFE
KSFSSTFRTLHLVNEIPVNERSHIPECSFKHDDSVAVDSLSSSDLQNQTNRIEHDLVNTVESQLVLFA
SDNQQLTHLRHSRSSPEADNRILNAIDRSNELKEFEIGLTMRKLQLKQSQLALSSHSHMLEKIKLSFG
FQKASFKGEKFKTRMQETRDAEILRTLIDFLVSAVIVMSACFGYGTYIYSYQRITDVTSACSATSKGS
KSWWMPNSVSNFSSGFLFLRCHVIAVTRMCFGILMILAIAWLAFQRSSTTGSNMPITFNLILLGIICG
FAGRFCTNTLGGDGNTWLMYWEVLCSIHLLGNLFPSLLYHVLHGPISVSHREQVVWLPYWVRRCLFYA
AVGLILPALTGLLPFASLSDWKDHFVEEIKSIVIGDKIEA*

**>oscpr5.1-B9**

MDAAAASSSSSSSATMAAAAASAAEASLSGSPASSRNARHRQRKGVRLRMLRRRGRQPV….QEDLALP
LGMSFAAVLAQVINTKNISGQRLHPDFLSKICTSAVKESLTNIYGDSSNSFIKNFEKSFSSTFRTLHL
VNEIPVNERSHIPECSFKHDDSVAVDSLSSSDLQNQTNRIEHDLVNTVESQLVLFASDNQQLTHLRHS
RSSPEADNRILNAIDRSNELKEFEIGLTMRKLQLKQSQLALSSHSHMLEKIKLSFGFQKASFKGEKFK
TRMQETRDAEILRTLIDFLVSAVIVMSACFGYGTYIYSYQRITDVTSACSATSKGSKSWWMPNSVSNF
SSGFLFLRCHVIAVTRMCFGILMILAIAWLAFQRSSTTGSNMPITFNLILLGIICGFAGRFCTNTLGG
DGNTWLMYWEVLCSIHLLGNLFPSLLYHVLHGPISVSHREQVVWLPYWVRRCLFYAAVGLILPALTGL
LPFASLSDWKDHFVEEIKSIVIGDKIEA*

**>oscpr5.1-B10**

MDAAAASSSSSSSATMAAAAASAAEASLSGSPASSRNARHRQRKGVRLRMLRRRGRQPVRGRARPGHG
GGGAVQEDLALPLGMSFAAVLAQVINTKNISGQRLHPDFLSKICTSAVKESLTNIYGDSSNSFIKNFE
KSFSSTFRTLHLVNEIPVNERSHIPECSFKHDDSVAVDSLSSSDLQNQTNRIEHDLVNTVESQLVLFA
SDNQQLTHLRHSRSSPEADNRILNAIDRSNELKEFEIGLTMRKLQLKQSQLALSSHSHMLEKIKLSFG
FQKASFKGEKFKTRMQETRDAEILRTLIDFLVSAVIVMSACFGYGTYIYSYQRITDVTSACSATSKGS
KSWWMPNSVSNFSSGFLFLRCHVIAVTRMCFGILMILAIAWLAFQRSSTTGSNMPITFNLILLGIICG
FAGRFCTNTLGGDGNTWLMYWEVLCSIHLLGNLFPSLLYHVLHGPISVSHREQVVWLPYWVRRCLFYA
AVGLILPALTGLLPFASLSDWKDHFVEEIKSIVIGDKIEA*

**>oscpr5.1-B11**

MDAAAASSSSSSSATMAAAAASAAEASLSGSPASSRNARHRQRKGVRLRMLRRRGRQPV……QEDLALP
LGMSFAAVLAQVINTKNISGQRLHPDFLSKICTSAVKESLTNIYGDSSNSFIKNFEKSFSSTFRTLHL
VNEIPVNERSHIPECSFKHDDSVAVDSLSSSDLQNQTNRIEHDLVNTVESQLVLFASDNQQLTHLRHS
RSSPEADNRILNAIDRSNELKEFEIGLTMRKLQLKQSQLALSSHSHMLEKIKLSFGFQKASFKGEKFK
TRMQETRDAEILRTLIDFLVSAVIVMSACFGYGTYIYSYQRITDVTSACSATSKGSKSWWMPNSVSNF
SSGFLFLRCHVIAVTRMCFGILMILAIAWLAFQRSSTTGSNMPITFNLILLGIICGFAGRFCTNTLGG
DGNTWLMYWEVLCSIHLLGNLFPSLLYHVLHGPISVSHREQVVWLPYWVRRCLFYAAVGLILPALTGL
LPFASLSDWKDHFVEEIKSIVIGDKIEA*

**>oscpr5.1-B12**

MDAAAASSSSSSSATMAAAAASAAEASLSGSPASSRNARHRQRKGVRLRMLRRRGRQPVGRARPGHGG
GGAVQEDLALPLGMSFAAVLAQVINTKNISGQRLHPDFLSKICTSAVKESLTNIYGDSSNSFIKNFEK
SFSSTFRTLHLVNEIPVNERSHIPECSFKHDDSVAVDSLSSSDLQNQTNRIEHDLVNTVESQLVLFAS
DNQQLTHLRHSRSSPEADNRILNAIDRSNELKEFEIGLTMRKLQLKQSQLALSSHSHMLEKIKLSFGF
QKASFKGEKFKTRMQETRDAEILRTLIDFLVSAVIVMSACFGYGTYIYSYQRITDVTSACSATSKGSK
SWWMPNSVSNFSSGFLFLRCHVIAVTRMCFGILMILAIAWLAFQRSSTTGSNMPITFNLILLGIICGF
AGRFCTNTLGGDGNTWLMYWEVLCSIHLLGNLFPSLLYHVLHGPISVSHREQVVWLPYWVRRCLFYAA
VGLILPALTGLLPFASLSDWKDHFVEEIKSIVIGDKIEA*

**Data S2**. Predicted amino acid sequence of *Os*CPR5.2 in wildtype and mutants. Amino acid sequences similar to the wildtype *Os*CPR5.1 sequence is noted in black, differences with the wildtype *Os*CPR5.1 sequence are noted in red.

**>*Os*CPR5.2**

MRSPPESGSRTGRGEEIAARQLNPTASASPPRSLIMDGACCDGGGSPESGGASSSASSYGSASRLQKG
VRLRRRRQRLRRPLLATGGDGRGAADGAQDLALPLGMSFAAVLAQVLNRSSCSEGRLQPDFLSKMCTS
AVKESLTNIYGDRFDNFTKNFEKSFGSTLRTLHLINETPVYEQDNSRFSHEDGTSAAEIKLSGADSKR
PVHDIQESTSLSSMDNQIILHAGTDQQLVKLPHNKASPEFDRHILNVFERSLNEQTRSNELKELEIGL
NMRKLQLKQSQIALSSYSHMLEKIKISMGFQKASFREEKFRTQMEDTRHAELLRRLIDLLLTAVVFMS
VCFGYGTYIYSYKRITAVTAACAAASREPKSWWMPNSVSAFNSGLLFFRCHLIAATRMSFGMLMILLI
AWLIFQRSAMTGPNMPITFNVMLLGVLCGSVGRFCVDTLGGDGNVWLFFWEILCFIHLFGNSRPSLLY
RMLYGPISVTDRTKASDLPYRVCRYTFYTVLSVILPCLAGLLPFASLSDWNELVVEYMKSKFIRINTE
V*

**>oscpr5.2-C1**

MRSPPEERVAHRSRRRNCGAATKPYRLCLAATLPYYGRGLLRRRWLAGVRRGVLVGVVVRLRVPAAEG
GAPAAAAAEAPETAACDWRGWEGRRRRRAGPRAASRDVLRRRFLPRSLIEAVALKEDYNLISFQRCAR
QQ*

**>oscpr5.2-C2**

MRSRAGRAPVEAKKLRRGN*

**>oscpr5.2-C3**

MSGSCPGP*

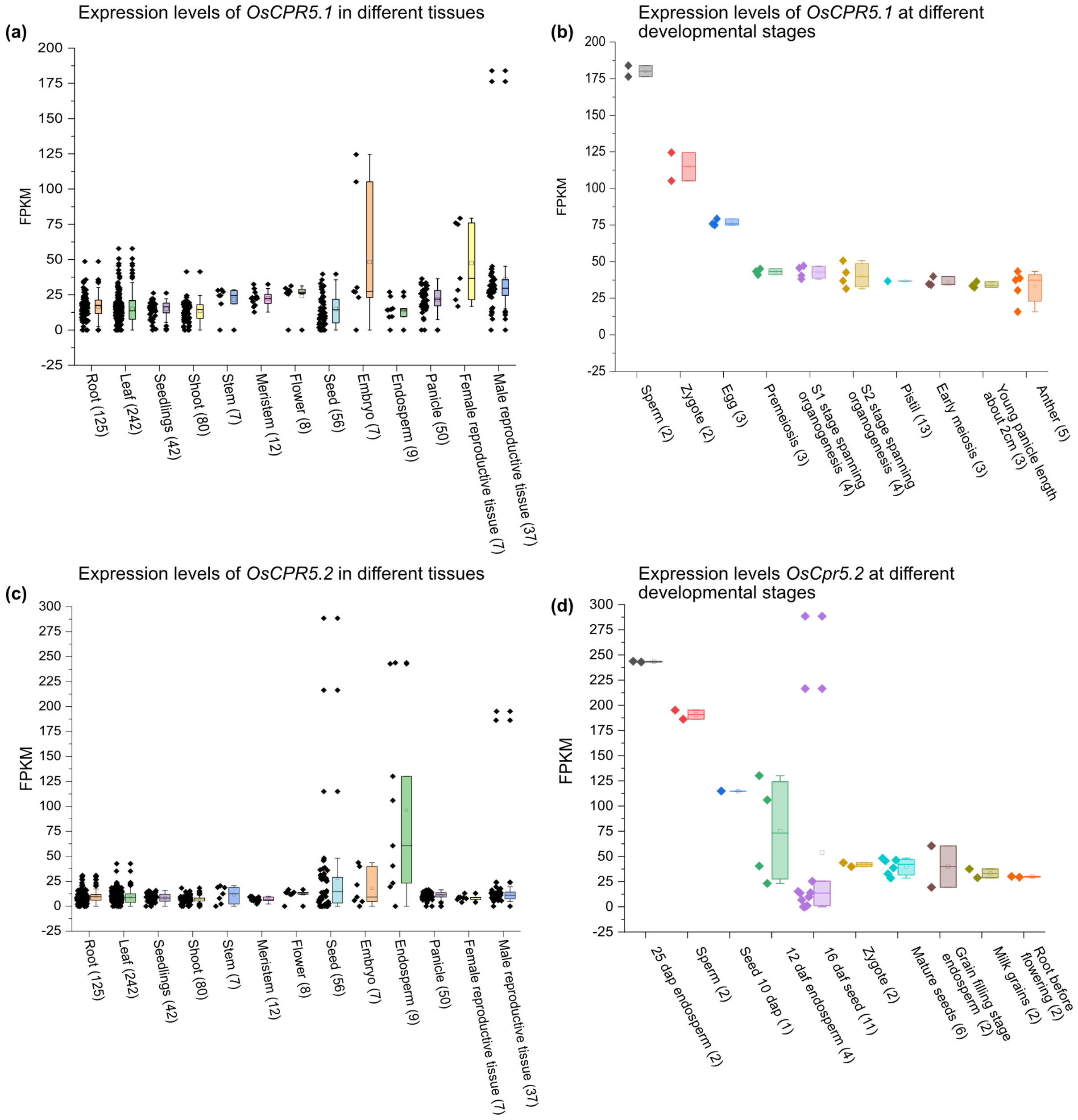


**Figure S1.** Tissue specific and developmental stages expression levels of *OsCPR5.1* and *OsCPR5.2*.

FPKM (Fragments Per Kilobase Million) data were retrieved from the Rice RNA Seq Database (Rice RNA-seq database, Zhailab@SUSTech) and analyzed using Origin software. Two genes showed broad and overlapping expression among different tissue and developmental stages and no major effects of various stresses on mRNA levels were found. Whiskers plot minimum and maximum values, asterisks indicate individual data points.

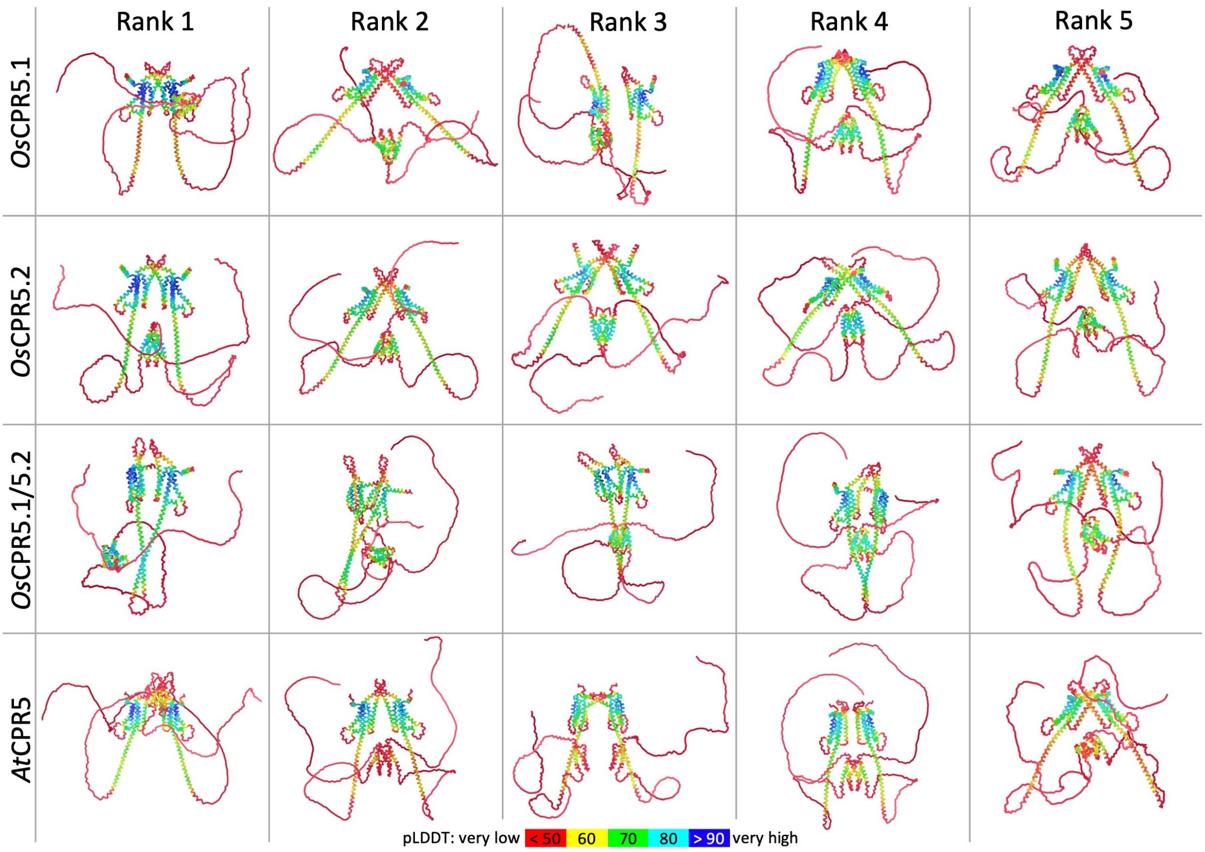


**Figure S2.** Alphafold prediction of *At*CPR5, *Os*CPR5.1 and *Os*CPR5.2 dimer conformations. Ranked dimer predictions of CPR5 from rice (*Os*) and Arabidopsis (*At*) were generated with the AlphaFold2* Colab tool using MMseqs2 and default settings. The color code indicate predicted local distance difference test (pLDDT) scores. *Os*CPR5.1 (*LOC_Os01g68970*), *Os*CPR5.2 (*LOC_Os02g53070*), *At*CPR5 (*At5g64930*). Note that prediction contains multiple domains marked in red with low pLDDT scores.

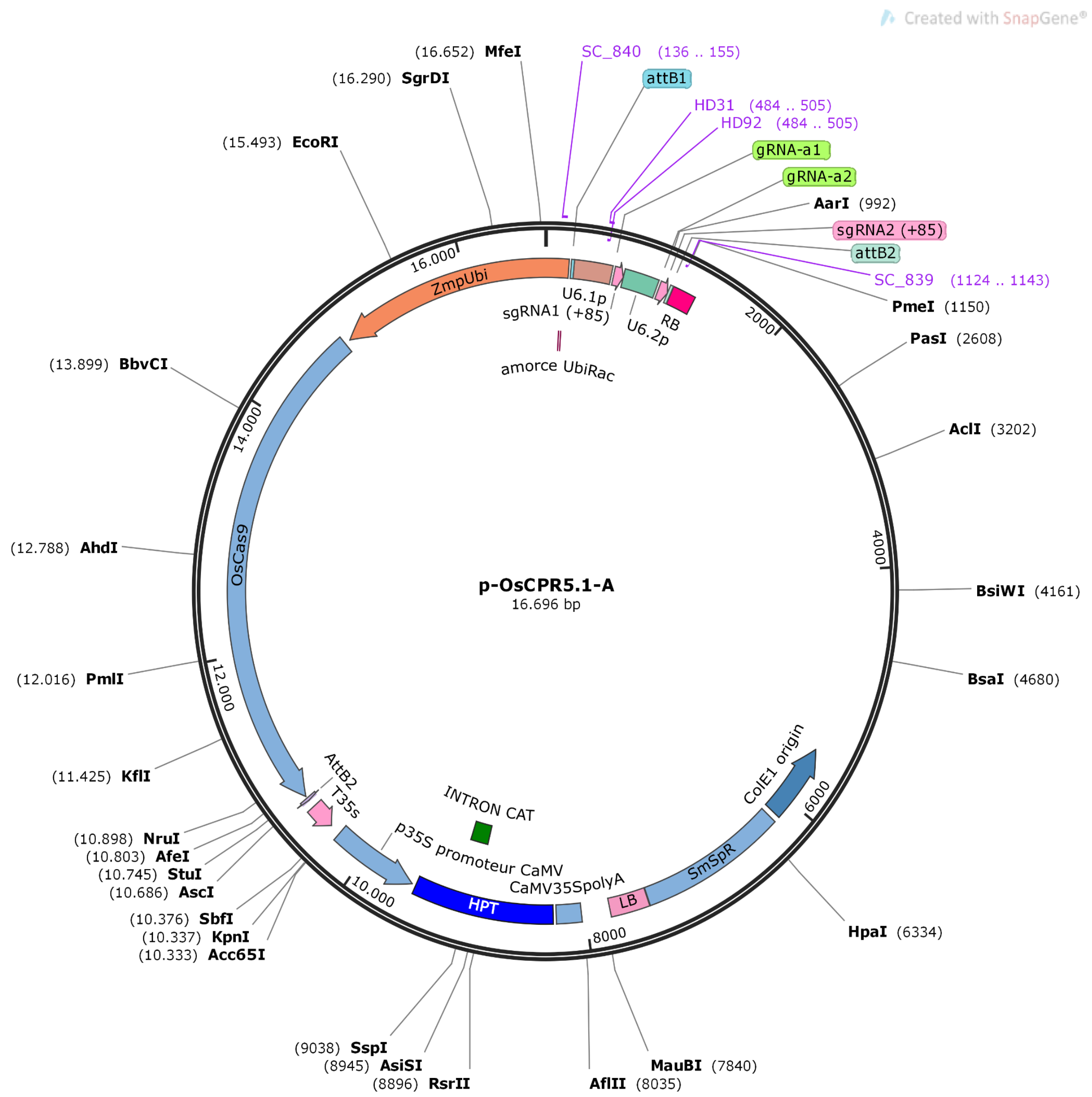


**Figure S3.** Map of the binary CRISPR/Cas9 vector p-OsCPR5.1-A.

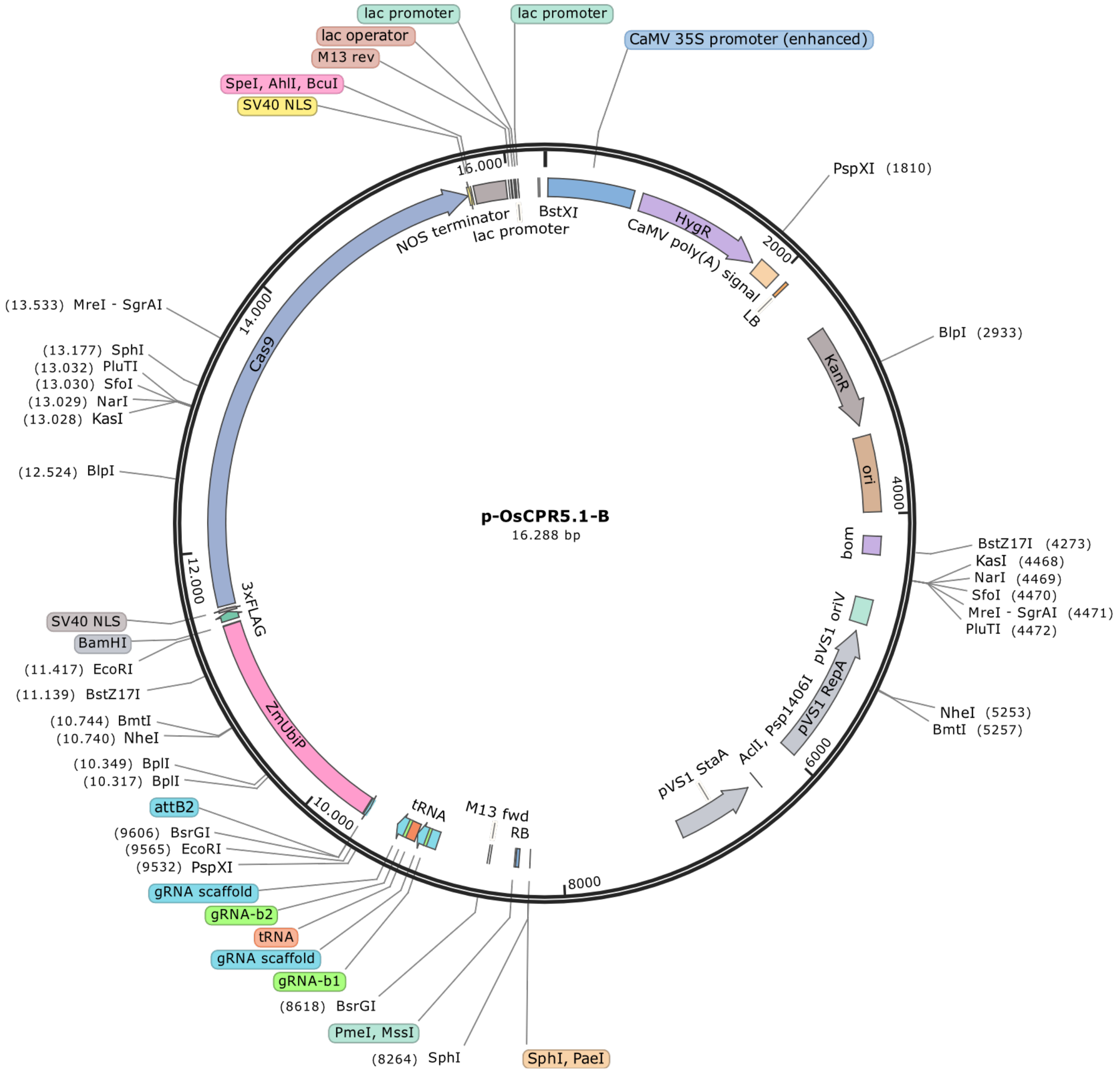


**Figure S4.** Map of the binary CRISPR/Cas9 vector p-OsCPR5.1-B.

(a)

(b)

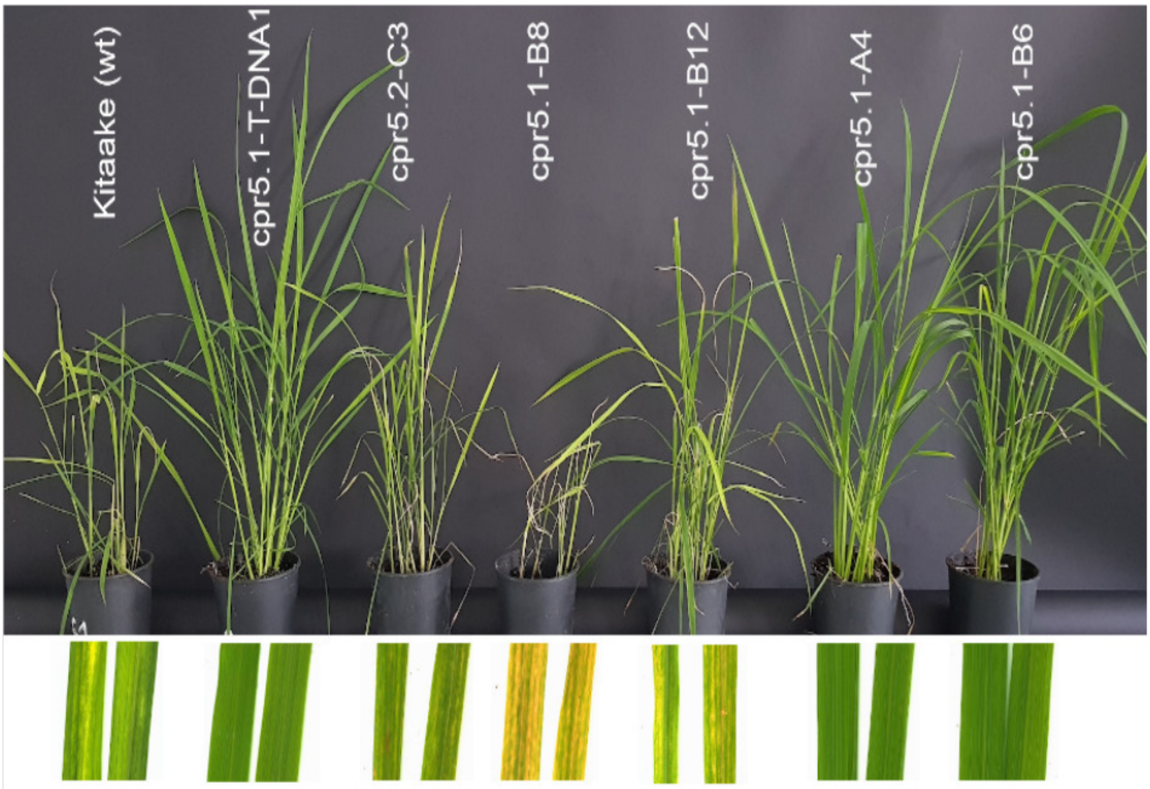


**Figure S5.** Disease resistance phenotypic reaction of *oscpr5.1* and *oscpr5.2* mutants. Plants were mechanically inoculated with BF1 isolate and were taken three weeks after inoculation. **(a)** Phenotype of controls (Kitaake wildtype and cpr5.1-T-DNA1) and mutants at IRD, plant age two weeks at time of inoculation. **(b)** Magnification of leaf area of RYMV-infected leaves of edited lines. Knockout mutants of *OsCPR5.1* (cpr5.1-A4, cpr5.1-B6) do not show obvious defects or symptoms. Knock-out mutant in *OsCPR5.2* (*oscpr5.2-C3*) and in frame mutants of *OsCPR5.1* (oscpr5.1-B8 and *oscpr5.1-B12*) show reduce size and leaves yellowing and mottling. Leaves symptoms appear to be more pronounced on in frame mutants, suggesting hypersensitivity to RYMV.

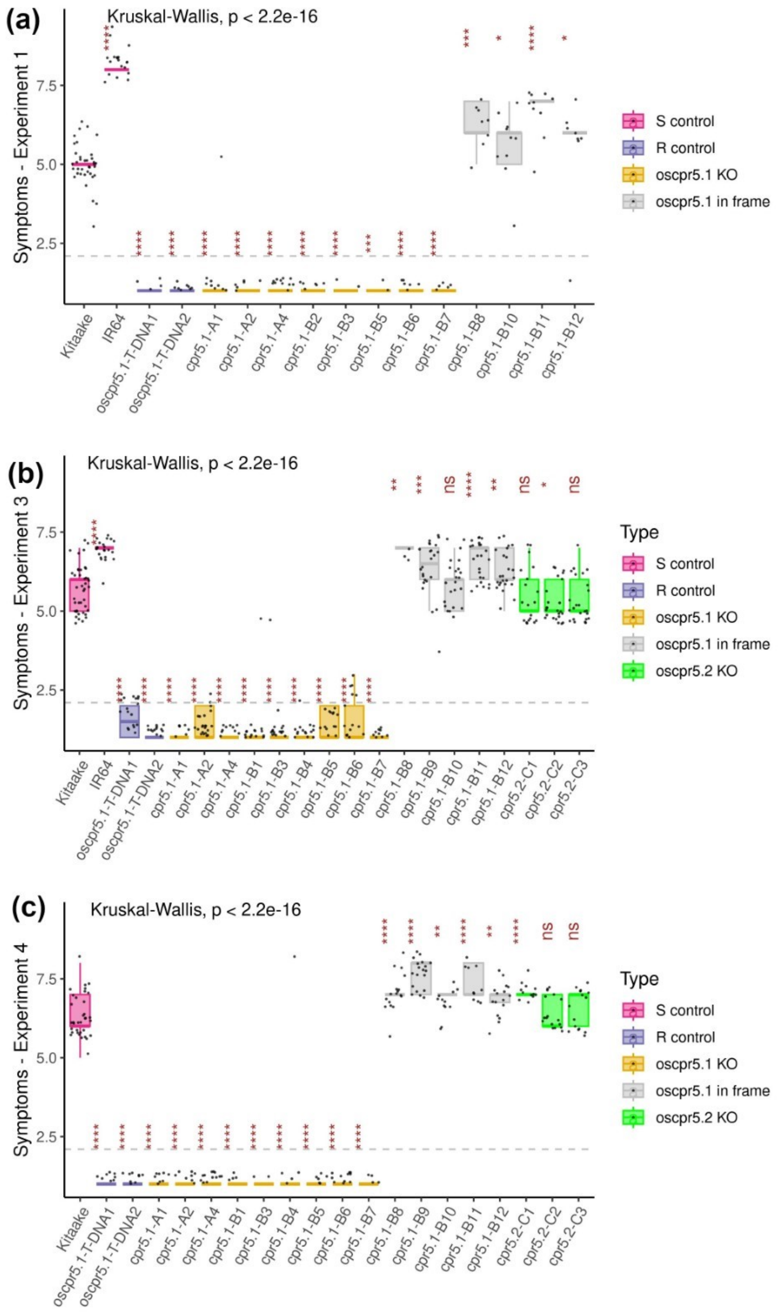


**Figure S6.** Symptoms of *oscpr5.1* and *oscpr5.2* mutant lines two weeks after inoculation with BF1 isolate of RYMV. Results of three independent experiments performed in greenhouse conditions are reported. Boxes extend from the upper (Q3) and lower (Q1) quartiles, the whiskers extend from Q3 +1,5 x the interquartile range (IQR) to Q1- 1,5 x IQR and the median values is represented by the center lines. Kruskall-Wallis test was used to detect significant differences between lines and each mutant lines was compared to Kitaake wildtype using pairwise Wilcoxon tests (ns: non-significant; *: p<0,05; **: p<0,01; ***: p<0,001; p<0,0001).

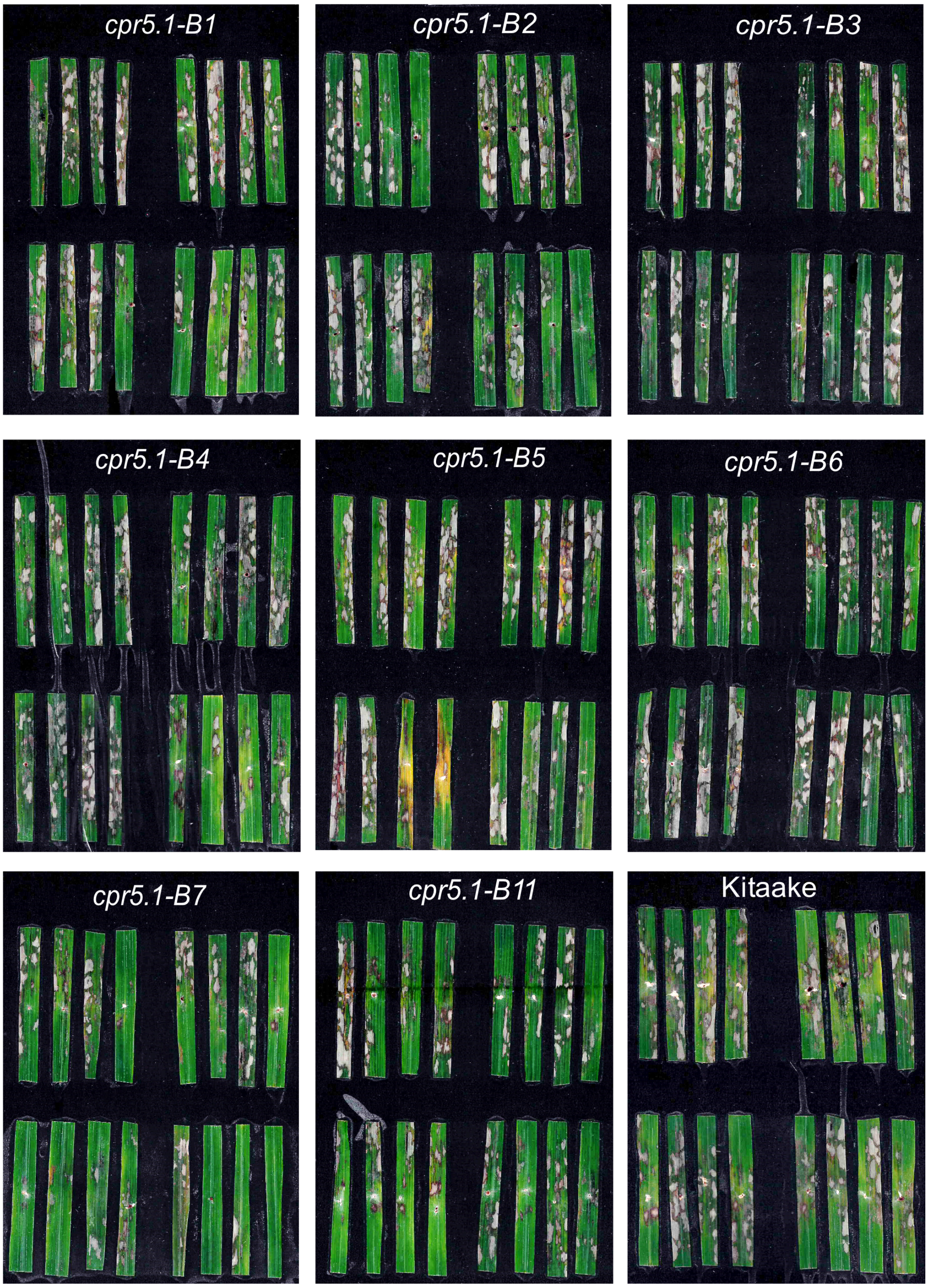


**Figure S7**. Disease phenotypic reaction of *oscpr5.1* mutants with *R. solani* AG1-1A. 30 days old plants were inoculated with *R. solani* (RSY-04) using the detached leaf assay method and observations were recorded 5 days of post inoculation. Single experiment.

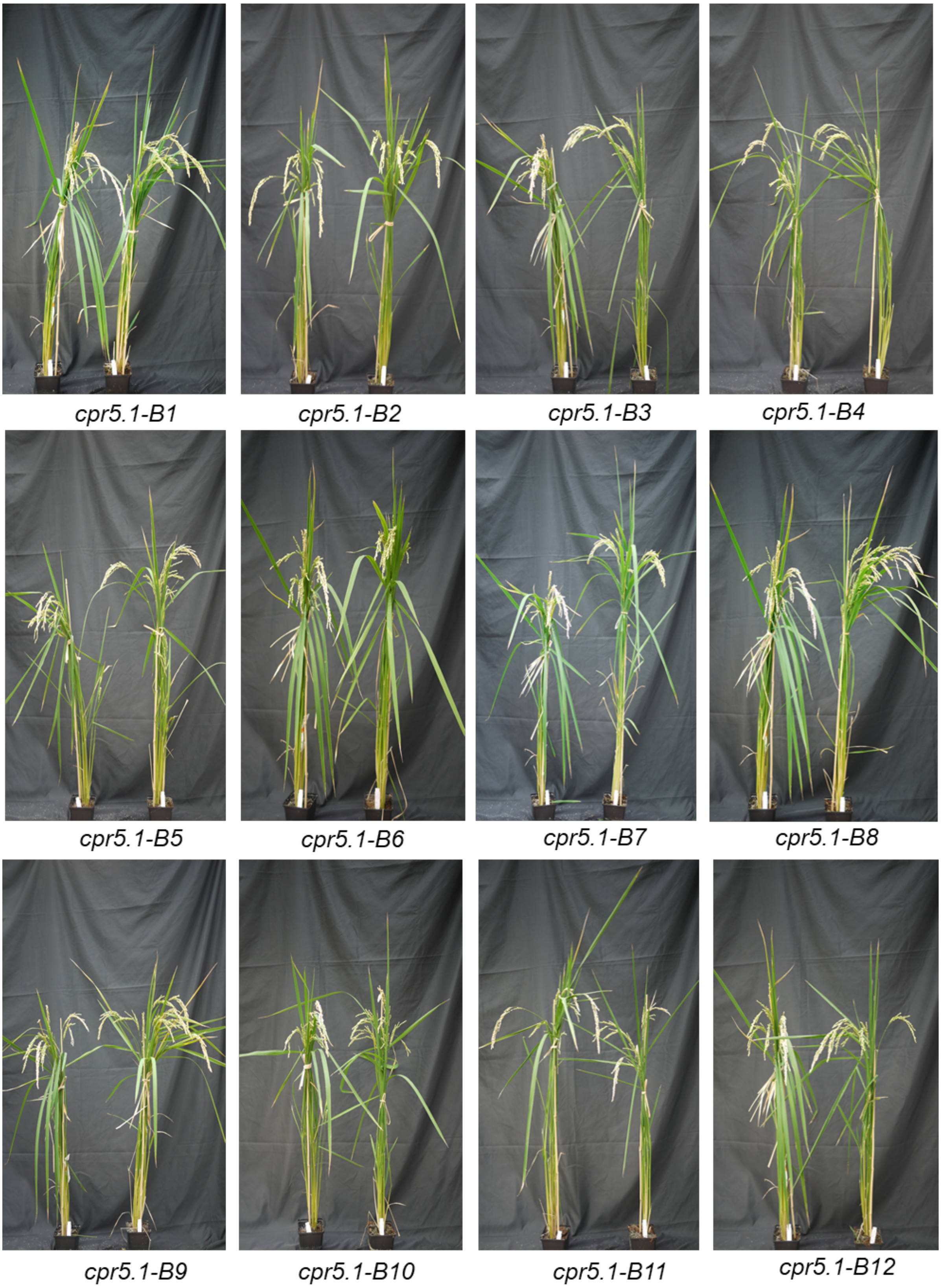


**Figure S8.** Phenotypic characters of *oscpr5.1* mutants. Left pot: Kitaake (wildtype), right pot: *oscpr5.1* mutant plant. Similar results were obtained in three independent experiments at HHU.

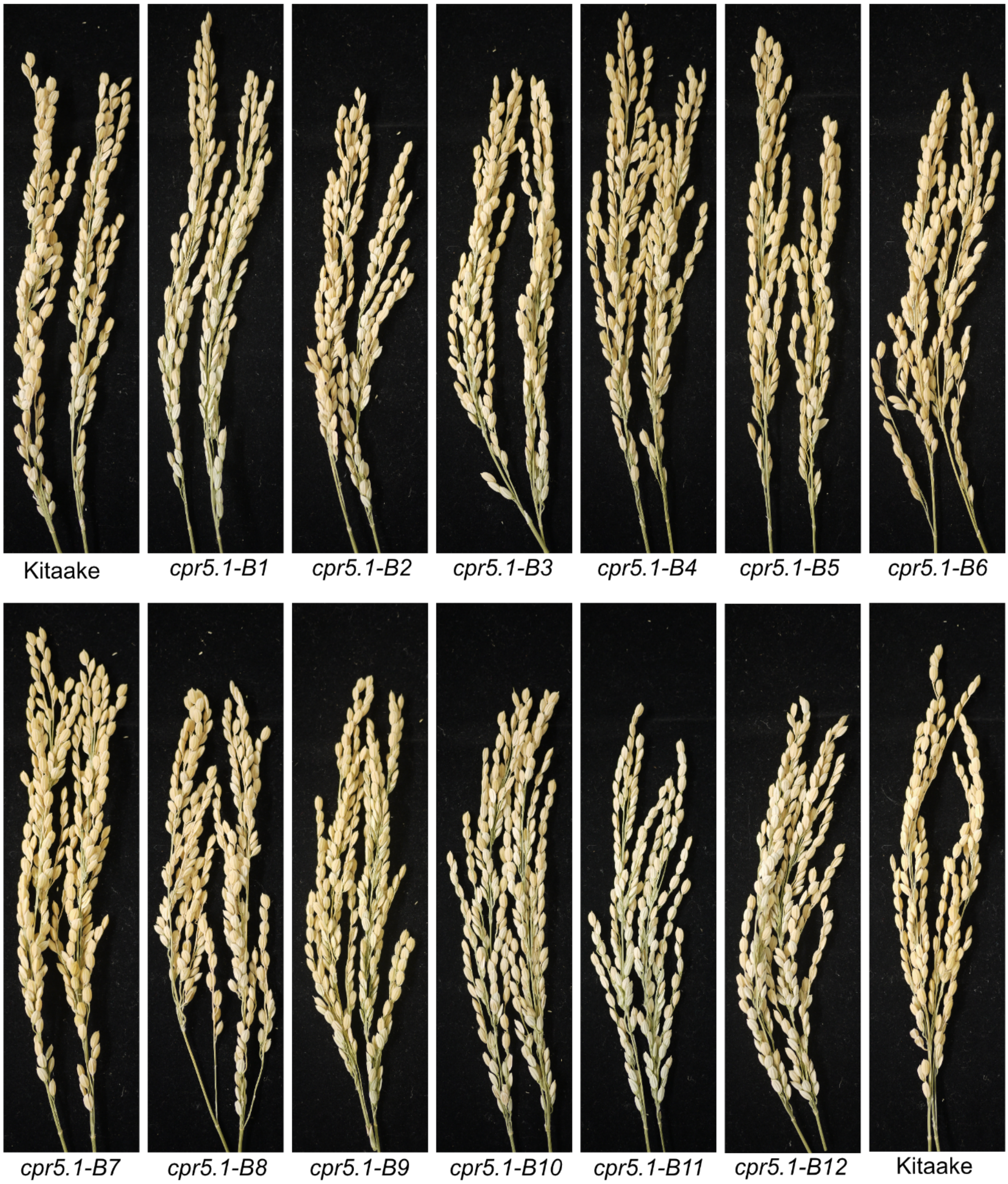


**Figure S9.** Panicle characters of *oscpr5.1* mutant plants. Comparable results were obtained in three independent experiments at HHU.

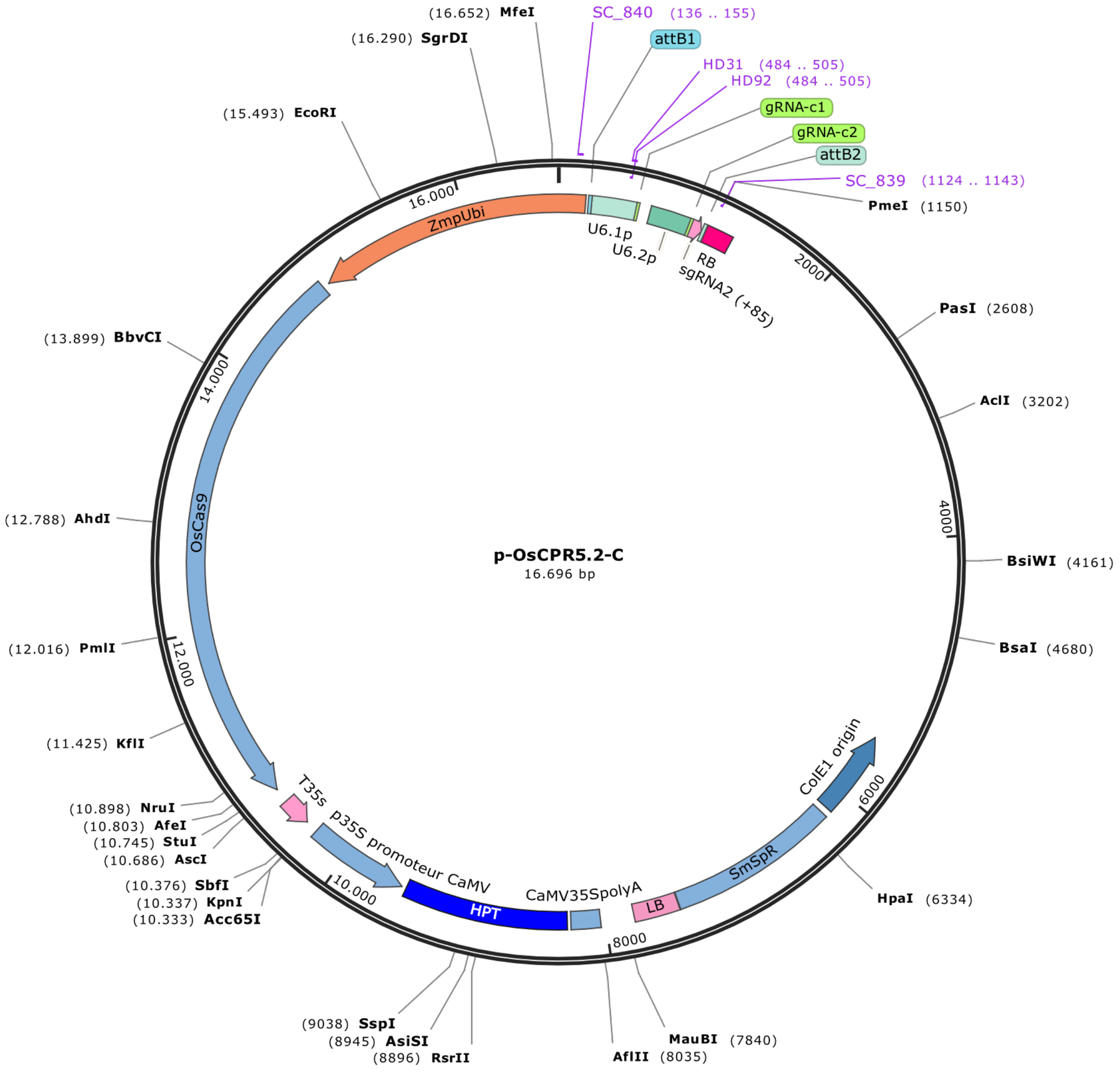


**Figure S10.** Map of the binary CRISPR/Cas9 vector p-OsCPR5.2-C.

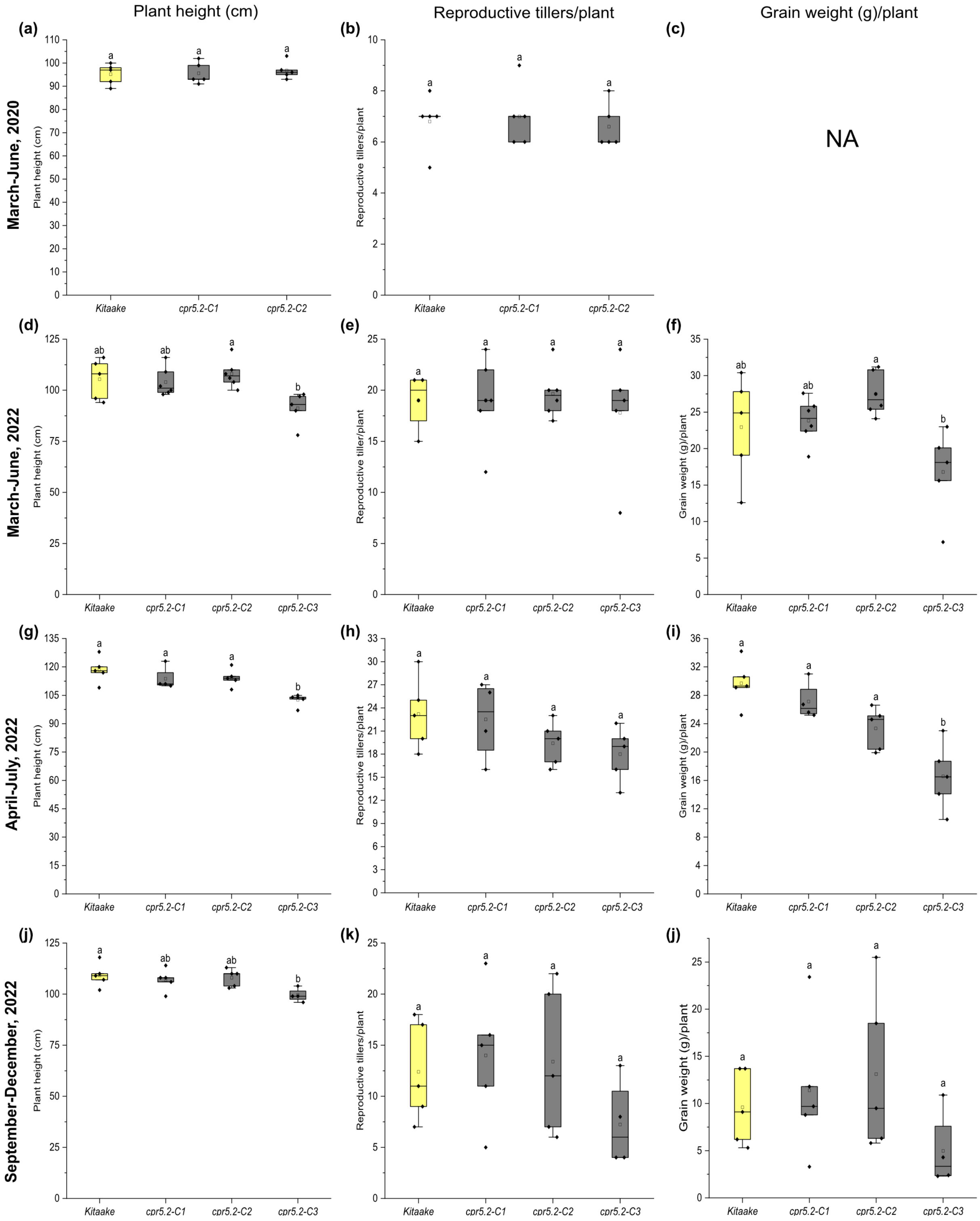


**Figure S11**. Morphological characters of *oscpr5.2* frameshift mutant plants. The data generated from four independent experiments conducted in greenhouses under controlled conditions. *cpr5.2-C3* was found to have reduced plant height (cm), reproductive tiller number/plant and grain weight (g)/plant. likely due to second site mutations. Boxes extend from 25th to 75th percentiles and display median values as center lines. Whiskers plot minimum and maximum values, asterisks indicate individual data points and detected by one-way ANOVA followed by Tukey's test, significance difference ($p< 0.05$). NA: not available.

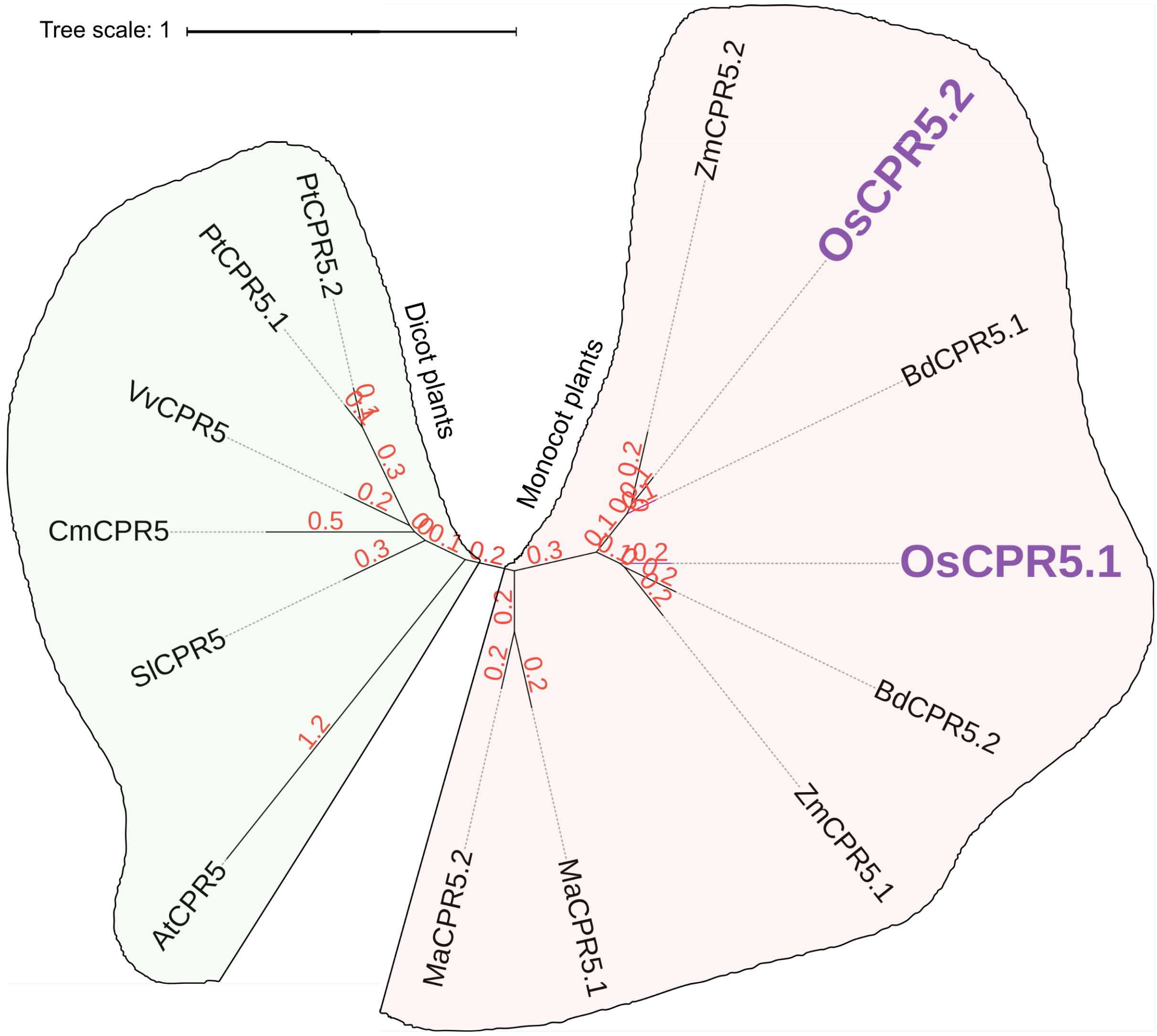


**Figure S12**. Alignment of CPR5 homologs compared to *Arabidopsis thaliana*. The unrooted phylogenetic tree was generated using the NGPhylogeny tool using the neighbor-joining method (https://ngphylogeny.fr) (Lemoine *et al.*, 2019) and visualized with the help of iTOL (https://itol.embl.de) (Letunic and Bork, 2019). Only the conserved regions of the protein sequences were considered for the analysis. Protein sequences were aligned using the MAFFT alignment program (Katoh *et al.*, 2019) with a gap-opening penalty of 1.53 and a gap-extension penalty of 0.123. The phylogenetic tree was generated using the neighbor-joining method and clade support scores were calculated by bootstrapping (n = 1.000), branch length values are displayed in red (number of substitutions per site). Dicot plants: *Arabidopsis thaliana* (At), *Cucumis melo* (cm), *Populus trichocarpa* (Pt), *Solanum lycopersicum* (Sl), *Vitis vinifera* (Vv). Monocot plants: *Brachypodium distachyon* (Bd), *Musa acuminata* (Ma), *Oryza sativa* (Os), *Zea mays* (Zm).

**Table S1.** List of gRNAs used to develop CRISPR/Cas9 mediated mutations of *OsCPR5.1* and *OsCPR5.2* in Kitaake

| **gRNA** | **Target sequence (5'-3') + PAM** | **Target region of CPR5 (downstream of start codon, ATG)** |
|---|---|---|
| CPR5.1-a1 | CGCCGCCATCGTCGCCGACG + **AGG** | 35 bp |
| CPR5.1-a2 | CGCCATGACGCACCTGCGCG + **AGG** | 267 bp |
| CPR5.1-b1 | GGCGCGCTCGGCCTCCAC + **AGG** | 175 bp |
| CPR5.1-b2 | CGAGCGCGCCCCGGGCGATG + **GGG** | 183 bp |
| CPR5.2-c1 | ATGAGAAGCCCACCGGAGAG + **CGG** | 1 bp |
| CPR5.2-c2 | TCTCGGGATGTCCTTCGCGG + **CGG** | 306 bp |

**Table S2.** List of primers used in the present study

| **Primer name** | **Sequence (5'-3')** | **Target for amplification** |
|---|---|---|
| CPR5.1-A1F(IRD) | CACGACGTCGCTTCGCCTCC | Primers for *OsCPR5.1* genotyping |
| CPR5.1-A1R(IRD) | TTTCCATTGAGGGAAGTATAGCA | |
| CPR5.1_B1F | GCGACACGAGGCTTTCTTC | |
| CPR5.1_B1R | GACGGTAAATGCCACACTACT | |
| CPR5.2-C1F (IRD) | TCCGAGTTTGCTTAACGGCT | Primers for *OsCPR*5.2 genotyping |
| CPR5.2-C1R (IRD) | GACGCCCAAAATGAGTTCCG | |
| M13-F | GTA AAA CGA CGG CCA GT | Primers for M13 |
| M13-R | CAG GAA ACA GCT ATG AC | |
| OsRAC-F | TCCATCTTGGCATCTCTCAG | Primers for RAC |
| OsRAC-R | GTACCCTCATCAGGCATCTG | |
| OsCas9-F | GGGTAATGAACTCGCTCTGC | Primers for Cas9 |
| OsCas9-R | TGGCGTCAAGAACTTCCTTTG | |
| Cas9-F(IRD) | TCACCTCCTTGTAGCCCTTG | |
| Cas9-R(IRD) | ACGGCGAGATTAGGAAGAGG | |
| Hygro-IIF | CCGCTCGTCTGGCTAAGATC | Primers for Hygromycin resistance |
| Hygro-IIR | GTCCTGCGGGTAAATAGCTG | |
| Hygro-F(IRD) | CTCGGAGGGCGAAGAATCTC | |
| Hygro-R(IRD) | GCTCCAGTCAATGACCGCTG | |

**Table S3:** List of CRISPR/Cas9-induced insertions or deletions in T2 homozygous plants of *OsCPR5.1*

| gRNA | Transgenic lines | Variations | Sequence | Mutation |
|---|---|---|---|---|
| gRNA-a1&a2 | *oscpr5.1-A1* | +1 bp, +1 bp | +T, +G | Frameshift |
| | *oscpr5.1-A2* | -232 bp | CGTCCTCCTCCTCCTCCTCC-GTGAGGTTATAAA | Frameshift |
| | *oscpr5.1-A4* | -45 bp, +1bp | ACGCCGCGGCGGCG-GCGGCCGCGGAGGCG, +G | Frameshift |
| gRNA-b1&b2 | *oscpr5.1-B1* | +1 bp, +1 bp | +A, +A | Frameshift |
| | *oscpr5.1-B2* | +1 bp, -1 bp | +T, -G | Frameshift |
| | *oscpr5.1-B3* | -40 | GGAGGCCGAGCGCGCCCCGGGCGATGGGGGCGGTGGCGCC | Frameshift |
| | *oscpr5.1-B4* | -3 bp, +1 bp | -GGA, +A | Frameshift |
| | *oscpr5.1-B5* | -26 | GGAGGCCGAGCGCGCCCCGGGCGATG | Frameshift |
| | *oscpr5.1-B6* | +1 bp, -50 bp | +T, CGGGCGATGGGGGCGGTGGCGCCGTGCAGGAGGATCTCGCGCTGCCTCT | Frameshift |
| | *oscpr5.1-B7* | -61 bp | GGCGGCAGCCTGTGGAGGCCGAGCGCGCCCCGGGCGATGGGGGCGGTGGCGCCGTGCAGGA | Frameshift |
| | *oscpr5.1-B8* | -1 bp, +1 bp | -G, +A | Substitution |
| | *oscpr5.1-B9* | -42 bp | GTGGAGGCCGAGCGCGCCCCGGGCGATGGGGGCGGTGGCGCC | In frame deletion |
| | *oscpr5.1-B10* | +1 bp, -1 bp | +A, -G | Substitution |
| | *oscpr5.1-B11* | -42 bp | GTGGAGGCCGAGCGCGCCCCGGGCGATGGGGGCGGTGGCGCC | In frame deletion |
| | *oscpr5.1-B12* | -2 bp, -1 bp | -GA, -G | Substitution |

* deletion: -; insertion + in number of base pairs, two numbers indicate biallelic state

**Table S5.** Virus detection using ELISA in symptomatic and symptomless plants at two weeks after inoculation with BF1 isolate of RYMV

| | **Experiment 1** | | **Experiment 2** | |
|---|---|---|---|---|
| | ELISA +** | ELISA - | ELISA + | ELISA - |
| Symptomatic* | 57 | 1 | 92 | 3 |
| Symptomless | 11 | 114 | 5 | 146 |

* Plants were considered as symptomatic when a disease score strictly higher than 2 was recorded

** ELISA tests were considered as positive when an optical density > 0.100 was obtained.

**Table S7:** List of CRISPR/Cas9-induced deletions in T2 homozygous plants of *oscpr5.2*.

| **gRNAs** | **Transgenic plants** | **Variations** | **Sequence** | **Mutation** |
|---|---|---|---|---|
| gRNA-c1&c2 | *oscpr5.2-C1* | +1 bp, +1 bp | +A, +T | Frameshift |
| | *oscpr5.2-C2* | -7bp, +1 bp | +CACCGGA, +C | stop |
| | *oscpr5.2-C3* | -320 bp | GAAGCCCACCGGAGAGCGGG<br>TCGCGCACCGGTCGAGGCGA<br>AGAAATTGCGGCGCGGCAAC<br>TAAACCCTACCGCCTCTGCCT<br>CGCCGCCACGCTCCCTTATTA<br>TGGACGGGGCCTGCTGCGAC<br>GGCGGTGGCTCGCCGGAGTCC<br>GGCGGGGCGTCCTCGTCGGCG<br>TCGTCGTACGGCTCCGCGTCC<br>CGGCTGCAGAAGGGGGTGCG<br>CCTGCGGCGGCGGCGGCAGA<br>GGCTCCGGAGACCGCTGCTTG<br>CGACTGGAGGGGATGGGAGG<br>GGCGCCGCCGACGGCGCGCA<br>GGACCTCGCGCTGCCTCTCGG<br>GATGTCCTTCGCG | Frameshift |

* deletion: -; insertion + in number of base pairs, two numbers indicate biallelic state
